# Expanded ACE2 dependencies of diverse SARS-like coronavirus receptor binding domains

**DOI:** 10.1101/2021.12.25.474149

**Authors:** Sarah M. Roelle, Nidhi Shukla, Anh T. Pham, Anna M. Bruchez, Kenneth A. Matreyek

**Affiliations:** Department of Pathology, Case Western Reserve University School of Medicine, Cleveland, OH, 44106, USA

## Abstract

Viral spillover from animal reservoirs can trigger public health crises and cripple the world economy. Knowing which viruses are primed for zoonotic transmission can focus surveillance efforts and mitigation strategies for future pandemics. Successful engagement of receptor protein orthologs is necessary during cross-species transmission. The clade 1 sarbecoviruses including SARS-CoV and SARS-CoV-2 enter cells via engagement of ACE2, while the receptor for clade 2 and clade 3 remains largely uncharacterized. We developed a mixed cell pseudotyped virus infection assay to determine whether various clade 2 and 3 sarbecovirus spike proteins can enter HEK 293T cells expressing human or *Rhinolophus* horseshoe bat ACE2 proteins. The receptor binding domains from BtKY72 and Khosta-2 used human ACE2 for entry, while BtKY72 and Khosta-1 exhibited widespread use of diverse rhinolophid ACE2s. A lysine at ACE2 position 31 appeared to be a major determinant of the inability of these RBDs to use a certain ACE2 sequence. The ACE2 protein from *R. alcyone* engaged all known clade 3 and clade 1 receptor binding domains. We observed little use of *Rhinolophus* ACE2 orthologs by the clade 2 viruses, supporting the likely use of a separate, unknown receptor. Our results suggest that clade 3 sarbecoviruses from Africa and Europe use *Rhinolophus* ACE2 for entry, and their spike proteins appear primed to contribute to zoonosis under the right conditions.

## Introduction

As shown by the ongoing severe acute respiratory syndrome-related coronavirus 2 (SARS-CoV-2) pandemic, viral spillover from animal reservoirs can decimate public health systems and the global economy. The likelihoods of zoonotic spillovers are multifactorial, including both ecological and molecular factors. Human disruptions to world ecosystems are increasing the likelihood of future zoonotic events[1]. We still lack a clear understanding of the molecular factors that likely play key roles during zoonosis.

Molecular compatibility during viral entry is a key determinant of viral tropism and host switching [2–6]. The *Betacoronavirus* genus include known zoonotic viruses of pandemic potential including Middle East Respiratory Syndrome-related coronavirus (MERS-CoV), SARS-CoV, and SARS-CoV-2. These viruses use the spike glycoprotein to catalyze entry into target cells upon binding to a compatible host cell receptor. Unlike MERS-CoV which uses Dipeptidyl-peptidase 4 (DPP4) as the cell surface receptor[7], the lineage B viruses of the sarbecovirus subgenus SARS-CoV and SARS-CoV-2 utilize angiotensin converting enzyme-2 (ACE2) as the host cell entry receptor[8, 9]. ACE2 binding from SARS-like CoVs is dictated by an independently folded domain of up to 223 residues in length, referred to as the receptor binding domain (RBD).

Multiple viral clades exist within the sarbecovirus subgenus, and the cell surface receptor dependencies of each clade are not well established[10, 11]. Clade 1 sarbecoviruses including SARS-CoV and SARS-CoV-2 are known to utilize ACE2, while the receptors for clade 2 and clade 3 viruses are unknown[10, 11]. The lack of observed ACE2-dependent enhancement to infection by clade 2 and clade 3 sarbecovirus spike proteins, such as YN2013 or BM48-31, can be explained in three ways: 1) these RBDs have weak but functionally relevant affinity for ACE2, below the limit of detection of commonly used assay, 2) these RBDs have affinity for certain orthologs of ACE2, but little or no affinity for human ACE2 or for any orthologs that have been tested so far, or 3) these RBDs primarily utilize an entry mechanism distinct from ACE2.

Here, we characterize the extent of ACE2 dependence across sarbecovirus clades. We utilized a single-copy HEK 293T genome modification platform to strongly overexpress multiple cell surface proteins proposed to serve as receptors for SARS-CoV-2, alongside the well-established receptor, ACE2 [12]. As the clade 2 and clade 3 sarbecoviruses were observed in samples collected from various *Rhinolophus* bats, we synthesized and expressed ACE2 orthologs from *R. ferrumequinum, R. affinis, R. alyone,* R. *landeri, R. pearsonii,* and various ACE2 alleles observed in *R. sinicus.* We observed differing patterns of ACE2 ortholog usage by various clade 3 sarbecoviruses RBDs during cell entry, including human ACE2-dependent entry by the BtKY72 and Khosta-2 RBDs. We observed little to no ACE2-dependent infection with RBDs from clade 2 sarbecoviruses, including various alleles from *R.sinicus* and *R.pearsonii* from which these viruses were isolated. Thus, our study provides a new genetic approach for characterizing receptor utilization during viral entry, and demonstrated that clade 3 sarbecoviruses likely utilize ACE2 as a cell-entry receptor during infection.

## Results

### Developing a robust genetic assay for viral entry

Knowing that sarbecovirus spike proteins may exhibit weak affinity for ACE2 proteins from mismatched hosts, we designed an assay for measuring biochemically weak but functionally important interactions promoting viral entry. We previously developed a Bxb1 recombinase-based transgenic expression system, wherein human ACE2 or its coding variants could be stably and precisely expressed by a Tet-inducible promoter already engineered into the cell genome, upon integration of a single promoterless plasmid[12] (**Fig 1A**). We found that the human ACE2 cDNA, when encoded behind a consensus Kozak sequence permitting frequent ribosomal translation of the mRNA, yielded high ACE2 cell surface abundance, roughly 10-fold greater than ACE2 protein observed in Vero-E6 cells, commonly used to propagate SARS-CoV or SARS-CoV-2 in cell culture[12].

**Figure 1.**
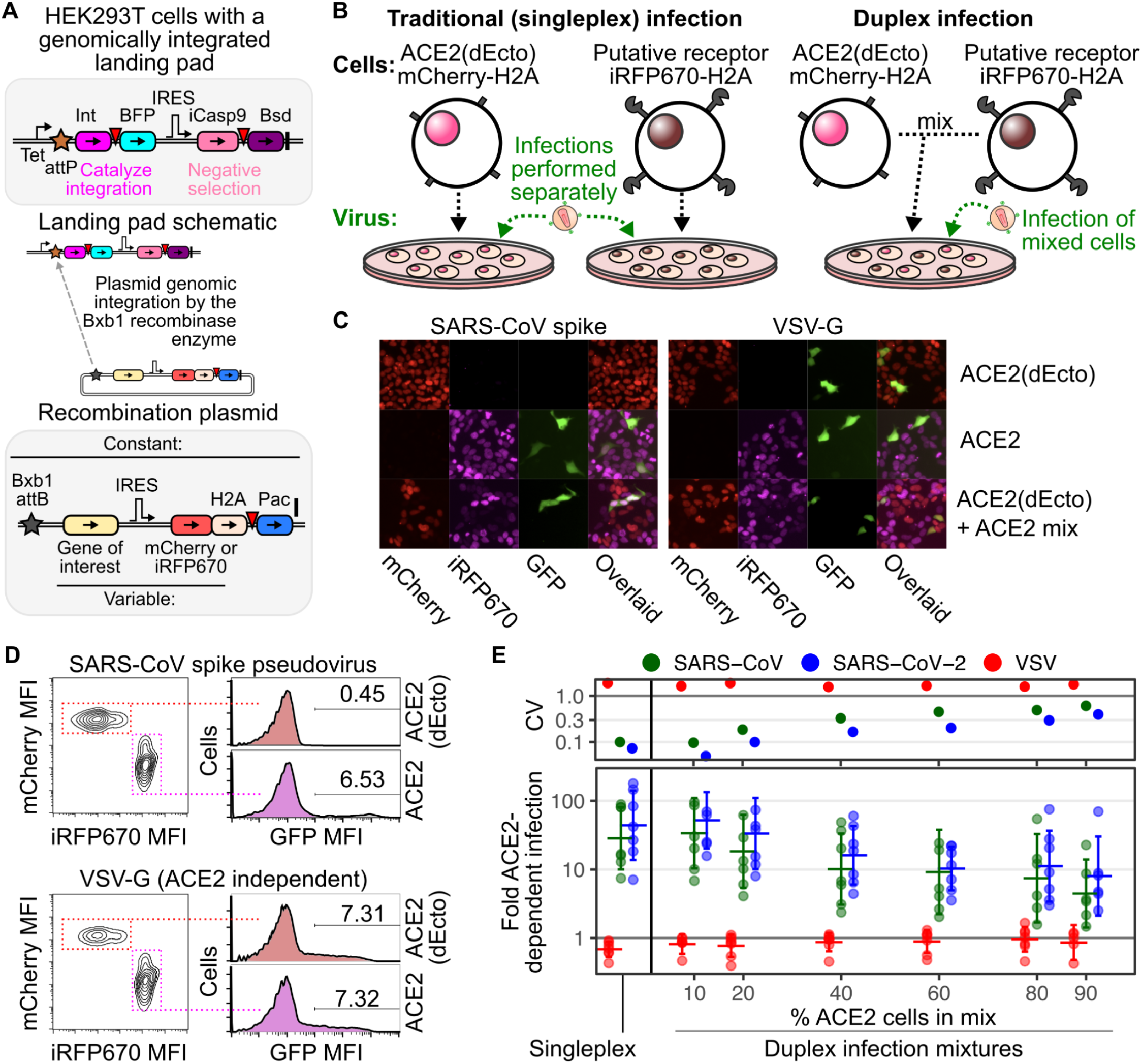
Duplex pseudovirus infection assay. A) Schematic of the Bxb1 recombinase “landing pad” engineered in HEK 293T cells, and the elements of the attB recombination vectors used to stably express transgenic DNA upon plasmid integration. Int, Bxb1 integrase; BFP, blue fluorescent protein; iCaspθ, inducible caspase 9; Bsd, blasticidin resistance gene; H2A, histone 2A; Pac, puromycin resistance gene. Inverted red triangles are 2A translational stop-start sequences. B) Cartoons describing the traditional singleplex and new duplex pseudovirus infection formats. C and D) Representative microscopy images (C) and flow cytometry profiles (D) of ACE2-dependent and independent pseudovirus infection in the duplex infection assay. E) Comparison of pseudovirus infection results obtained using the singleplex and duplex formats. Error bars denote 95% confidence intervals from 7 replicate experiments. Fold increase to infection in ACE2 overexpressing cells over non-expressing cells is shown on the bottom, while the coefficient of variation (CV) of the replicate results are shown on the top.

We previously discovered that our pseudovirus infection system was more sensitive than traditional *in vitro* binding assays utilizing soluble proteins. For example, expression of ACE2 mutants K31D or K353D, which reduced binding to soluble monomeric SARS-CoV RBD *in vitro[13]*, had little to no effect for SARS-CoV spike pseudovirus infection when translated from a consensus Kozak sequence in our expression platform[12]. Instead, we only observed reduced pseudovirus infection when the K31D or K353D ACE2 mutant protein levels were reduced 30-fold, suggesting that avidity effects conferred by high cell-surface ACE2 abundances can compensate for reductions to binding affinity. Thus, we focused on further developing a flexible pseudovirus infection assay capable of detecting weak but specific protein interactions enabling infection.

To increase throughput, we converted the traditional singleplex pseudovirus infection format into a duplex assay configuration. Traditionally, control and experimental cells are plated separately into different wells (**Fig 1B, left**). All wells are then exposed to the same volume of viral inoculum and infectivity is quantitated by taking a ratio of the amount of infection present in the experimental wells divided by the amount of infection present in the control wells. While ensemble measurements such as luciferase activity require a traditional singleplex format, fluorescent reporters for infection, such as GFP positivity, are single-cell assays and easier to multiplex. Thus, we developed an approach wherein the experimental and control cells are marked by different fluorescent proteins, allowing the two cell types to be mixed together and infected by the same inoculum of GFP-reporter pseudovirus within the same well (**Fig 1B, right**). Instead of calculating the ratio of GFP positivity from two different wells, we take the ratio of GFP positivity in mCherry negative control cells or putative receptor overexpressing iRFP670 positive cells, all from a single well. When testing two nearly isogenic cell lines differing solely by their expression of a putative receptor transgene, this ratio quantifies the amount of receptor-dependent enhancement to infection that has occurred.

To validate this approach, we created ACE2(dEcto) negative control HEK 293T cells encoding human ACE2 lacking its entire ectodomain, and thus incapable of serving as a cell surface receptor for SARS-CoV or SARS-CoV-2 spike. We marked these cells with red nuclei using mCherry-fused histone H2A (**Fig 1B**). Notably, HEK 293T cells naturally express a low but detectable amount of endogenous ACE2 from the X chromosome[12], thus accounting for the low, background level of infection in the assay. We next created ACE2 HEK 293T cells encoding full-length human ACE2 and marked these cells with near-infrared fluorescent nuclei using iRFP670-fused histone H2A. These cells exhibit more than 100-fold increased ACE2 protein than unmodified HEK 293T cells[12]. These cells were mixed into the same well and exposed to GFP encoding lentiviral particles coated with the ACE2-dependent envelope glycoprotein SARS-CoV spike (**Fig 1C, left**), or an ACE2-independent envelope glycoprotein such as vesicular stomatitis virus glycoprotein (VSV-G; **Fig 1C, right**), which uses LDLR as the viral entry receptor[14]. After two or more days, the entire well of cells can be analyzed with multi-color flow cytometry to simultaneously measure the infection rates in ACE2-expressing or control cells. We observed that ACE2-dependent viruses, such as those with SARS-CoV spike, exhibited preferential infection of the ACE2 expressing, iRFP670 fluorescent cells, whereas pseudoviruses coated with VSV-G infected the mCherry and iRFP670 expressing cells equally (**Fig 1D**). We will heretofore refer to this as the duplex infection assay.

We next performed a systematic analysis of how the duplex infection assay performed when the two cells were mixed at different ratios and compared these results with data obtained using the traditional singleplex assay format. We observed the greatest ACE2-dependent infection when the ACE2-expressing cells were a tenth of the total cells in the well (**Fig 1E**), with the coefficient of variation similar to the traditional singleplex assay format. As the proportion of ACE2-expressing cells increased, the amount of ACE2-dependent SARS-CoV-2 spike mediated infection reduced from ~ 52-fold at 10% ACE2-expressing cells to ~ 16-fold at 40% ACE2-expressing cells. There was a concomitant increase in the coefficient of variation suggesting a loss of data precision, at least partially due to insufficient sampling of the background level of infection in the control cells. Thus, when the receptor-expressing cells were infrequent in the mixed pool of cells, the mixed-cell infection assay was capable of producing data of comparable magnitude and precision to the traditional singleplex assay format, while reducing the number of total samples and requiring fewer physical manipulations. Due to these largely favorable characteristics, we used the duplex infection assay for all subsequent experiments.

### Known and proposed receptors for SARS-CoV-2 spike-mediated infection

While ACE2 is the primary SARS-CoV-2 spike receptor, numerous other proteins have been suggested to serve as alternative receptors. *BSG* encodes CD147 / Basigin, which was proposed to be a novel host cell receptor for SARS-CoV-2[15], though this has since been refuted[16, 17]. *CLEC4M* encodes L-SIGN / CD209L, which along with the related DCSIGN / CD209, was proposed to be a receptor[18, 19], and glycomimetic antagonists can block this interaction and inhibit SARS-CoV-2 infection[20]. Lectins are generally regarded as attachment factors rather than *bona fide* receptors as they have been implicated in enhancing entry for over 30 different viral glycoproteins[21]. *NRP1* and *NRP2* encode Neuropilin-1 / NRP1 and Neuropilin-2 / NRP2, which were proposed to be receptors since they bind peptides formed upon furin cleavage[22, 23], and the SARS-CoV-2 spike has a furin cleavage site that is important for its transmission[24], while SARS-CoV spike does not.

We used our duplex infection assay platform to compare the functional impacts of these proposed alternative receptors with ACE2 during SARS-CoV and SARS-CoV-2 spike pseudotyped virus infection. We created plasmid constructs encoding both untagged and cytoplasmically HA-tagged cDNAs of each proposed receptor protein (**Fig 2A**), and genomically integrated these DNAs alongside an IRES-iRFP670-H2A cassette into HEK 293T cells. Cells encoding the HA-tagged versions were immunoblotted to confirm the expression of each protein (**Fig 2B**). The predominant bands in the ACE2, CD147, and L-SIGN lysates corresponded to the electrophoretic migration sizes of the full-length, glycosylated proteins. In contrast to these three proteins, the bands corresponding to NRP1 and NRP2 were less abundant, although bands consistent with full-length protein were seen upon longer exposure (**Fig 2B, top**). Thus, all five proteins were expressed from their corresponding cDNAs, albeit to varying steady-state abundances.

**Figure 2.**
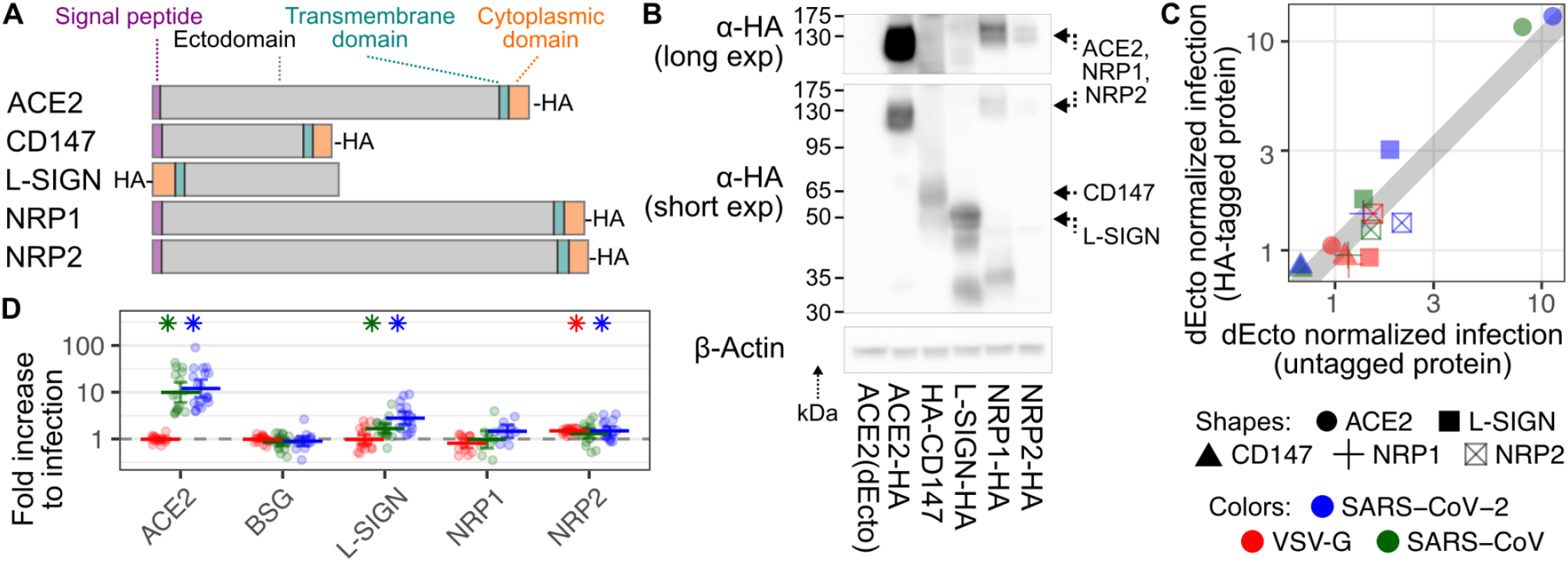
Established and proposed SARS-CoV-2 receptor proteins. A) Schematic showing relative lengths, tag locations, and overall domain topologies of proteins. B) Representative immunoblot of cells stably expressing HA-tagged versions of each protein, including longer (top) and shorter (bottom) anti-HA exposures. Beta actin was blotted as a loading control. C) Scatter plot comparing normalized pseudovirus infection rates of cells stably expressing various HA-tagged and untagged proteins. Shapes correspond to receptors. Colors correspond to viruses. D) Compiled normalized infection data for the established and proposed SARS-CoV-2 receptor proteins. Error bars denote 95% confidence intervals for at least 10 replicates. Asterisks denote samples with p < 0.01.

We next determined how the presence of each protein enhanced SARS-CoV and SARS-CoV-2 entry. Pseudovirus infection of cells expressing the HA-tagged or untagged forms of the protein were nearly identical (Pearson’s r^2^: 0.98, n = 15; **Fig 2C**). Thus, to improve statistical power for weak effect sizes, we merged the two datasets. Consistent with the known importance of ACE2, its expression increased infection with SARS-CoV 10-fold and SARS-CoV-2 spike 12-fold, while pseudoviruses with VSV-G were unaffected. L-SIGN increased SARS-CoV-2 spike infection 2.8-fold, while it had a more modest 1.7-fold effect on SARS-CoV spike (**Fig 2D**). The next strongest effect was a 1.5-fold increase to SARS-CoV-2 spike mediated infection conferred by NRP2, though it similarly simulated infection by VSV-G, which is not processed by furin. Thus, our assay revealed that ectopic ACE2 expression conferred the strongest increase to SARS-CoV and SARS-CoV-2 spike-mediated pseudovirus infection while L-SIGN conferred a milder but still significant increase.

### The BtKY72 RBD confers human ACE2-dependent infection

Following the initial SARS-CoV outbreak of 2002, there were extensive efforts to identify the bat viruses that were its precursor. This resulted in the isolation of WIV1, a bat virus highly related to SARS-CoV capable of using human ACE2 for entry [25]. Similar surveillance efforts uncovered hundreds of related coronaviruses in bats, but many of the receptor usages of these viruses are unknown. Due to the increased sensitivity possible with pseudovirus assays[12], we focused our remaining studies toward assessing the ACE2 dependencies of diverse, uncharacterized SARS-like CoV spike proteins.

To further establish the specificity of the approach, we subjected the mixture of cells expressing either the human ACE2 cDNA or the ectodomain-deleted control ACE2 construct to a panel of viruses pseudotyped with a wide range of viral entry glycoproteins. We found that the glycoproteins for Ebolavirus, Marburgvirus, Lassa fever virus, Lymphocytic choriomeningitis virus (LCMV), Junin virus, and Middle Eastern Respiratory Syndrome coronavirus (MERS-CoV) infected both ACE2 expressing and ACE2 null cells similarly (**Fig 3A**), consistent with the fact that none of these glycoproteins rely on ACE2 for infection [14, 26–29]. In contrast, the spike protein from WIV1 exhibited clear human ACE2-dependence comparable to SARS-CoV and SARS-CoV-2 spike (**Fig 3A**). Thus, the ACE2 duplex pseudovirus infection assay can be used to query the dependencies of a wide range of viral glycoproteins with high specificity.

**Figure 3.**
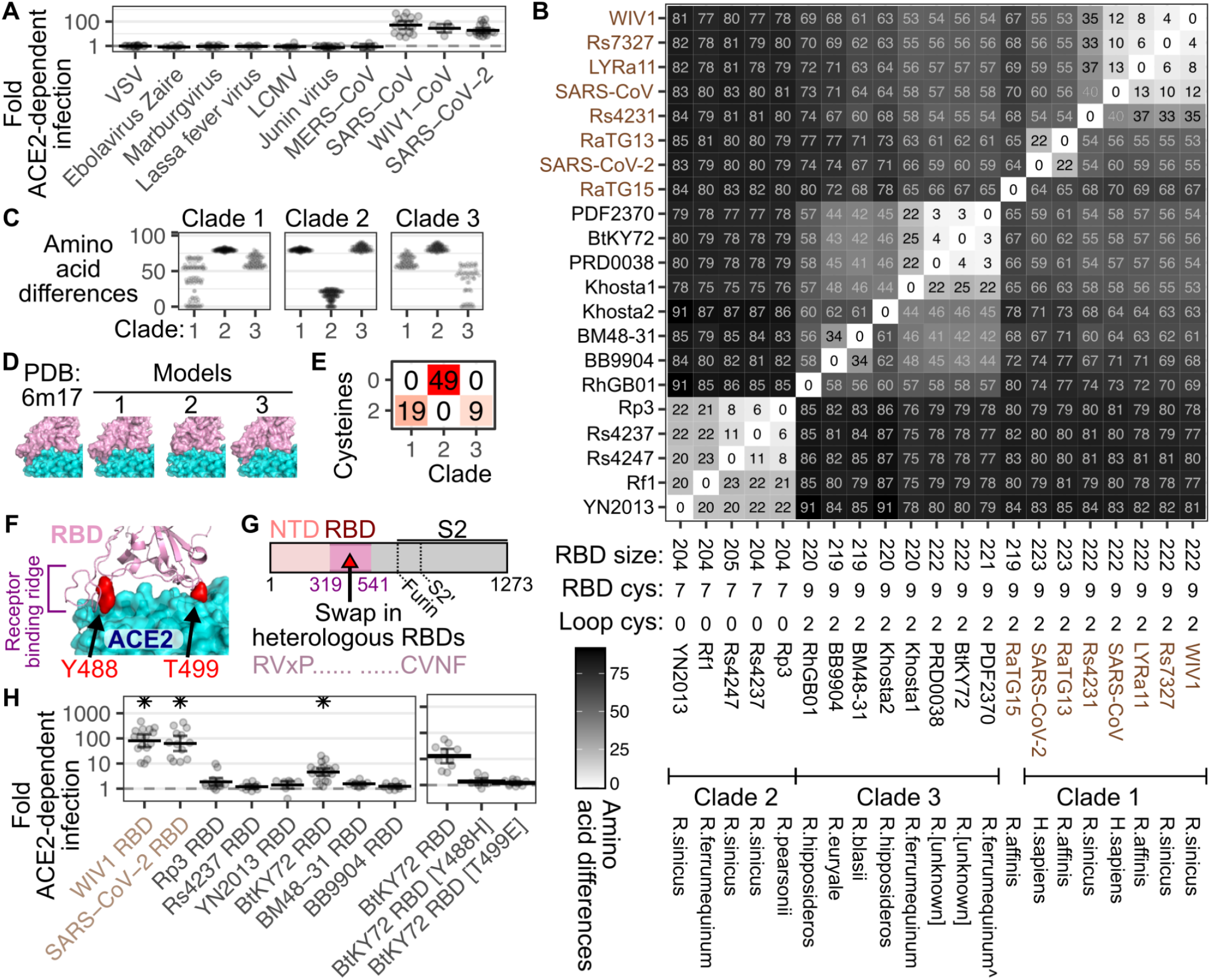
Diversity in viral glycoprotein sequences. A) Normalized human ACE2 dependent infection data from 5 or more replicate pseudotyped virus infections with diverse viral entry glycoproteins. B) Hamming distance matrix between the pairs of RBD amino acid sequences. Numbers denote non-identical amino acids between each pair. RBD lengths, virus names, clade groupings, and bat host species identified upon isolation are shown at the bottom. Names in brown were previously shown to bind human ACE2. The caret symbol denotes the closest bat species inferred through ACE2 protein sequencing from the sample. C) Non-identical residues between RBDs grouped by clade. D) Models of RBD tertiary structure, with the RBD colored pink and ACE2 colored in cyan. SARS-CoV RBD sequence was used to make the model for clade 1, YN2013 RBD sequence for clade 2, and the BtKY72 RBD sequence for clade 3. E) Number of RBDs with cysteines found within amino acids 158 through 171, separated by clade. F) Cartoon representation of the BtKY72 RBD model, denoting the receptor binding ridge and the residues mutated in panel G shown as red spheres. G) Cartoon schematic denoting the overall constructions of the transgenic RBD SARS-CoV chimeric spikes. H) Normalized infection data with chimeric spike proteins (left), with subsequent BtKY72 RBD mutant spike infection results shown (right). Asterisks denote p < 0.01. Error bars denote 95% confidence intervals.

We next turned our attention to characterizing the RBDs from the spike proteins of novel sarbecoviruses observed in bats. The RBDs from the spike proteins of sequenced sarbecoviruses thus far divide into three major clades [10, 11], although more are likely to be found and divided into additional clades[30]. All RBDs from clade 1 viruses tested so far have used ACE2 for entry, oftentimes exhibiting clear binding and utilization of human ACE2 [10], while the receptor dependencies of clade 2 and clade 3 viruses are unknown [10, 11]. We compiled a list of clade 2 and clade 3 RBDs following manual curation of sarbecovirus spike proteins currently listed in the National Center for Biotechnology Information (NCBI) (**Supplementary Table 1**). Clade 3 RBDs were all roughly 219 to 222 residues in length, and thus similar in length to clade 1 RBDs which were between 222 and 223 residues. In contrast, clade 2 RBDs possess an internal deletion, resulting in RBDs of 204 or 205 residues in length. To contextualize the protein sequence differences between diverse sarbecovirus RBDs, we created a matrix of pairwise Hamming distances of the differences in amino acid sequence for each RBD, ordered by hierarchical clustering (**Figs 3B and SFig 1)**. The resulting clustering recreated the three established clades [10] and largely corresponded to evolutionary phylogenies [11].

Clade 1 and clade 3 virus RBDs exhibited similarities not shared with clade 2 RBDs. Clade 1 and clade 3 RBD sequences were more similar to each other than clade 2 RBDs, while clade 2 RBDs were equally distant from both of the other clades (**Figs 3B and 3C**). Clade 1 RBDs also exhibited high intra-clade variability, such as between the SARS-CoV and SARS-CoV-2 RBDs, which exhibit 60 amino acid differences (**Fig 3B**). Similar to clade 1 RBDs, the clade 3 RBDs exhibited high intra-clade variability, with many pairwise combinations differing by 40 or more amino acids (**Figs 3B and 3C**). In contrast, the RBDs from clade 2 viruses yielded comparatively low intra-clade variability, with the RBDs exhibiting 28 amino acid differences or less (**Figs 3B and SFig 1**). In a previous study, none of the twenty one clade 2 RBDs tested used human ACE2[10]. Only three clade 3 RBDs (BM48-31, PRD-0038, and PDF-2386) were tested, with none exhibiting increased infection with human ACE2 [10, 11]. In contrast, RaTG15, which possesses an RBD distinct from the other known sarbecoviruses and thus may constitute a separate clade[30], and the clade 1 viruses, have all been shown to use ACE2, either from humans or from other animals.

The dissimilarity of the clade 2 RBD sequences relative to the other clades likely results in an altered tertiary structure in the RBD surface typically known to bind ACE2. Homology modeling of the three-dimensional structure of the YN2013 clade 2 RBD showed the ~15 residue deletion to cause a shortening of the receptor binding motif in a section often referred to as the “receptor binding ridge”[31](**Fig 3D**), a disulfide-linked loop that makes contact with the N-terminal alpha helix in the ACE2 protein ectodomain (**Fig 3F**). This deletion removes the disulfide bond, as none of the clade 2 RBDs encode cysteines in that region (RBD residues 158 through 171), while all clade 1 or clade 3 viruses do (**Fig 3E**). Accordingly, there are only seven cysteines encoded in clade 2 RBDs, while the clade 1 and clade 3 viruses have nine (**Fig 3B)**. In contrast, homology models of the BtKY72 clade 3 and SARS-CoV-2 clade 1 RBDs showed an extended receptor binding ridge similar to the experimentally determined SARS-CoV-2 ACE2 cryo-EM co-structure [31] (**Fig 3D)**.

Coronavirus spike proteins are routinely greater than 1250 amino acids in length, and chemically synthesizing each cDNA for functional analysis is prohibitively expensive. We instead took a chimeric spike approach, where we chemically synthesized the RBDs from various bat coronaviruses, and inserted the sequence in place of the RBD of SARS-CoV spike (**Fig 3G**) [10, 11]. We tested a panel of three clade 2 RBDs: Rs4237 [32], Rp3 [33], and YN2013 [34], each found in horseshoe bats in Eastern or Southeastern Asia. We also initially tested a small panel of clade 3 RBDs: BM48-31 [35], BB9904, and BtKY72 [36], which were the only three clade 3 RBDs identified as of June 2020. BM48-31 and BB9904 were identified from *R. blasii* and *R. euryale* bat samples collected in Bulgaria, while BtKY72 was collected from an unidentified *Rhinolophus* bat in Kenya. As positive controls, we generated chimeric SARS-CoV spikes encoding the WIV1 or SARS-CoV-2 RBDs. The WIV1 and SARS-CoV-2 RBDs promoted strong ACE2 -dependent pseudovirus entry, corresponding to 81-fold and 63-fold, respectively (**Fig 3H, left**). Of the clade 2 and clade 3 RBDs, only BtKY72 exhibited significant human ACE2-dependent entry, corresponding to a ~ 5-fold increase.

To validate our result with BtKY72, we explored this interaction in additional contexts. We first assessed whether we could enhance ACE2-dependent entry by co-expressing the cell surface protease TMPRSS2, which promotes spike-mediated viral entry at the cell surface [37–39], bypassing the need for endocytosis and proteolysis by endosomal cathepsins for activation [40]. The BtKY72 chimeric virus exhibited slightly increased infection in the presence of TMPRSS2 (**SFig 2A**). Furthermore, upon alignment of the BtKY72 RBD with the SARS-CoV, WIV1, and SARS-CoV-2 RBDs, we found a number of highly conserved positions at the purported ACE2 interface, including BtKY72 residues Y488 and T499 (corresponding to Y489 and T500 in SARS-CoV-2) (**Fig 3F**). To test whether these residues were involved in BtKY72 RBD interaction with human ACE2, we created Y488H or T499E mutant BtKY72 RBD chimeric spike pseudoviruses. Both single amino acid mutations abrogated ACE2-dependent entry (**Fig 3H, right**). While our observation of human ACE2 dependence with the BtKY72 RBD chimeric SARS-CoV spike should be reflective of what happens with the full-length BtKY72 spike, this needs to be formally shown. We attempted to recreate the full BtKY72 spike by piece-wise addition of additional BtKY72 spike sequence into the BtKY72 RBD chimeric spike, but were unable to observe ACE2 dependent entry (**SFig 2B**), as additional sequence swap points resulted in largely non-functional protein.

### Multiple clade 3 sarbecoviruses use human and rhinolophid ACE2

Having observed human ACE2 utilization by the RBD from BtKY72, we undertook a more comprehensive experiment testing combinations of diverse RBDs with various horseshoe bat or human ACE2 proteins. During our previous experiments, six more clade 3 RBDs were identified. Three were identified through the USAID-PREDICT project, corresponding to PRD-0038, PDF-2370, and PDF-2386. Like BtKY72, these sequences were discovered in African bats of unknown species within the *Rhinolophus* genus [11]. The RBDs from PDF-2370 and PDF-2386 were identical, and all three RBDs clustered closely with BtKY72 (**Fig 3B**), so we did not incorporate them into our study. In contrast, three highly unique RBDs were also discovered through ecological sequencing efforts; Khosta-1 and Khosta-2, observed in *R. ferrumequinum* and *R. hipposideros* bats in Sochi Russia [41], and RhGB01 observed in a *R. hipposideros* bat in the United Kingdom [42]. We thus created SARS-CoV chimeric spikes encoding these RBDs.

Based on our aforementioned result with BtKY72, we suspected that more clade 3 sarbecovirus RBDs are ACE2-dependent, but may only be compatible with ACE2 sequences encoded by their natural hosts, prompting us to synthesize additional ACE2 orthologs from *Rhinolophus* bats. ACE2 is known to be positively selected in bats, particularly at residues at the interface with SARS-CoV spike [43]. ACE2 is also highly polymorphic in *R. sinicus* bats [43, 44]. Due to these variations, different horseshoe bat ACE2 sequences likely exhibit a range of compatibility with diverse RBD sequences. There are more than 106 *Rhinolophus* species known [45]. The majority of the variation between rhinolophid ACE2 sequences are found in the ACE2 ectodomain, including positions 24, 27, 31, and 34, which exhibited 4 or more different amino acids at each site (**Fig 4A**). These highly variable positions are along the face of an alpha helix in contact with SARS-like CoV RBDs including their receptor binding ridge (**Fig 4B**), and were previously shown to be positively selected [43].

**Figure 4.**
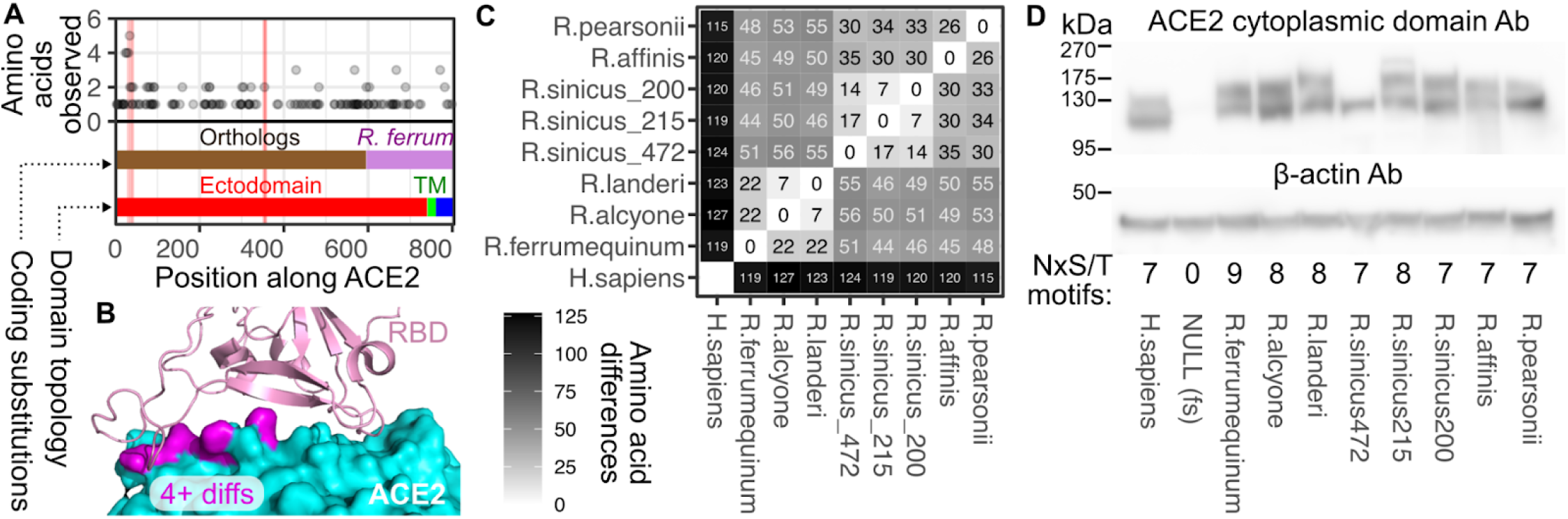
Diversity in ACE2 protein sequences. A}ACE2 position with the greatest number of amino acid differences amongst horseshoe bats. The red vertical lines denote the two segments of ACE2 linear sequence involved in its interaction with SARS-CoV and SARS-CoV-2 RBDs: positions 31 through 41 for the first patch, and 353 through 357 for the second. The ACE2 coding regions from the panel of horseshoe bats are shown in dark brown, while the C-terminal portion common to all *Rhinolophus* ACE2 constructs is colored purple. The ectodomain, transmembrane, and cytoplasmic domains are colored in red, green, and blue, respectively. B) Positions with four or more different amino acids observed across rhinolophid ACE2 proteins as seen in panel A are shown in magenta in a structure of SARS-GoV-2 RBD bound to human ACE2 (PDB; 6m17). C) Hamming distance matrix showing the number of non-īdentical amino acids between pairs of tested ACE2 chimeric proteins. D) Representative immuno bl Otting of the total steady-stale abundance of each chimeric ACE2 protein stably expressed, alongside a 2A-1 inked open reading frame encoding human TMPRΞΞ2, in HEK 293T landing pad cells. An antibody raised against the human ACE2 cytoplasmic sequences, but also cross-reactive with the sequence present in R. *ferrumequinum.* was used in the top panel. An antibody that recognizes human beta-actin was used in the bottom panel. The number of potential N-glycosylation motifs in each sequence are shown at the bottom.

To quantitate the differences between the ACE2 sequences, we created another Hamming distance matrix comparing each pair of rhinolophid ACE2 protein sequences (**SFig 3**). We then chose 6 orthologs, including 3 distantly related alleles from *R. sinicus*, for experimental characterization. We synthesized codon-optimized cDNA sequence encoding the first 600 ACE2 residues, corresponding to the majority of the ectodomain of each *Rhinolophus* ACE2, as this region should contain all of the sequence impacting RBD binding and viral entry. To minimize effort and cost, we ligated these sequences to a DNA sequence encoding the last 200 residues of *R. ferrumequinum* ACE2, including its transmembrane and cytoplasmic regions (**Fig 4A**). A distance matrix for our tested constructs showed that the rhinolophid ACE2 sequences differed by a minimum of 7 residues and a maximum of 56 (**Fig 4C**). Each chimeric ACE2 allele was stably recombined into HEK 293T landing pad cells, and expression was confirmed by immunoblot against the *R. ferrumequinum* cytoplasmic domain, shared by all of our chimeric rhinolophid ACE2 constructs (**Fig 4D**). All constructs yielded ACE2 proteins at roughly the expected size, with slight differences in electrophoretic mobility among samples, potentially due to differing numbers of N-glycosylation motifs. Instead of a doublet like the rest of the samples, the *R. sinicus* (*472*) allele yielded a single band in some immunoblot experiments (**Fig 4D**), although it also yielded a doublet in other replicate immunoblots.

We performed the duplex pseudotyped virus infection assay as a matrix of combinations of chimeric RBD pseudotyped viruses and cells expressing human or *Rhinolophus* ACE2 proteins. The fraction of control mCherry+ or ACE2-expressing miRFP670+ cells were quantitated using flow cytometry and averaged across the replicates (**Fig 5A**). In the process of performing these experiments, we observed massive syncytia formation in a subset of samples (**SFig 4**), including some syncytia that reached half a millimeter in width (**Fig 5B)**. Since large syncytia are unlikely to be efficiently measured alongside normal sized cells in the flow cytometer, we also quantified the frequency of green fluorescence in mCherry+ or miRFP670+ cells using fluorescent microscopy (**SFig 5**). Despite the reduced dynamic range of the microscopy assay, the results between the two measurements were largely consistent (**Figs 5D and SFig 6**), providing additional confidence in the results. Regardless of the readout, the controls also performed as expected, wherein VSV-G coated pseudoviruses did not exhibit ACE2-dependent infection and miRFP670+ cells expressing a null ACE2 cDNA harboring an early frameshift mutation did not enhance spike-mediated pseudovirus infection above that of mCherry+ ACE2(dEcto) cells (**Figs 5A and 5C**).

**Figure 5.**
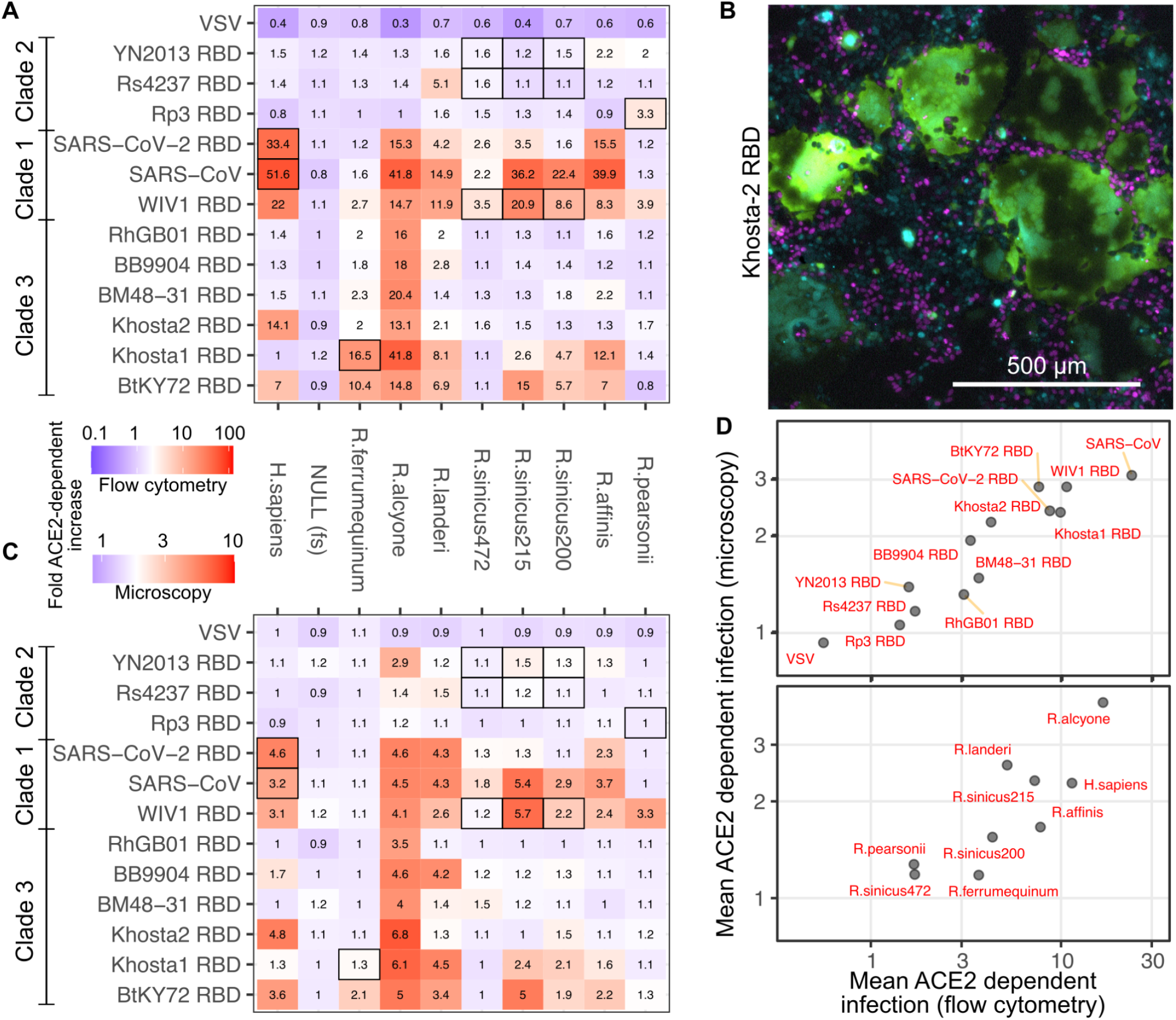
Infection of cells expressing rhînolophid ACE2 ectodomains with RBD chimeric spike protein pseudoviruses. A) Heatmap showing the amount of ACE2-dependent infection observed between various combinations of ACE2 proteins and RBD chimeric SARS-CoV spike proteins, or VSV-G, as measured with flow cytometry. Inset numbers denote the geometric means of replicate experiments. Black box outlines denote the host species recorded at time of virus sampling and sequencing. B) Representative image of large GFP+ syncytia observed in *R. alcyone* ACE2 cells infected with Khosta-2 RBD pseudovirus, and a subset of other samples. Magenta and cyan shapes are mCherry+ and iRFP670+ nuclei, respectively. C) To avoid disruption of syncytia, the amount of ACE2-dependent infection was quantified by looking for mCherry or iRFP670 fluorescence overlap with GFP fluorescence in microscopic images of adherent cells. D) Scatter plots showing the mean ACE2-dependent infectivities for each RBD observed across all cell lines expressing various ACE2 orthologs (top), or the mean ACE2-dependent infectivilies for each ACE2 ortholog across all RBDs tested (bottom). All constructs encode human TMPRSS2 C-terminally linked to ACE2 with a 2A translational stop-start sequence.

The clade 1 RBDs showed broad utilization of ACE2 alleles (**Figs 5A, 5C and 5D**), typified by WIV1, which was clearly enhanced by all but two orthologs tested. This included strong enhancement by *R. sinicus(215)*, consistent with the fact that WIV1 was isolated from *R. sinicus* bats of currently unknown allelic genotype. Also in agreement with previous reports, neither SARS-CoV nor SARS-CoV-2 RBDs utilized ACE2 from *R. ferrumequinum* or *R. pearsonii* [46], while both utilized ACE2 from *R. affinis[47, 48]* and *R. alcyone[9]*. SARS-CoV RBD exhibited compatibility with ACE2 from *R. sinicus,* much like WIV1, consistent with this likely being similar to its initial source of zoonosis. In contrast, SARS-CoV-2 RBD exhibited low compatibility with the tested *R. sinicus* alleles, but did exhibit strong compatibility with ACE2 from *R. affinis.* This is consistent with RaTG13, isolated from *R. affinis* bats, possessing one of the most similar RBD sequences to SARS-CoV-2 to date[49, 50].

In contrast, clade 2 RBDs did not exhibit obvious signals of ACE2-dependent entry (**Figs 5A and 5C**). Only the Rs4237 RBD paired with *R. landeri* ACE2 yielded signal in both the flow cytometry and microscopy readouts (**Figs 5A and 5C**), although this effect was relatively minor. We also did not observe consistent signals with alleles belonging to the *R. sinicus* bat species that the viruses were originally isolated from (**Figs 5A and 5C, black boxes**). As a whole, pseudoviruses generated with clade 2 RBD chimeric spike proteins exhibited an intermediate level of ACE2 dependence between the other sarbecovirus RBDs and VSV-G (**Fig 5D, top**), suggesting that there may still be slight ACE2 binding and utilization over background. Regardless, the lack of strong compatibility between the tested clade 2 RBDs and any of the *Rhinolophus* ACE2 alleles we tested suggest that these viruses use a different cell surface protein as a primary receptor and at best, only use ACE2 in an auxiliary role.

Clade 3 RBDs exhibited several distinct patterns of ACE2 ortholog usage. Along with BtKY72, the Khosta-2 RBD exhibited clearly enhanced entry in the presence of human ACE2. These two RBDs differed by 46 amino acids (**Fig 3B**), corresponding to ~ 80% amino acid identity, suggesting that they may be using a different set of amino acid side chain interactions to engage the ACE2 protein surface. Accordingly, their usage patterns of the tested *Rhinolophus* alleles were vastly different. Khosta-2 could only utilize ACE2 from *R. alycone,* while BtKY72 exhibited the broadest utilization of ACE2 alleles in the panel. The pattern was largely similar to the clade 1 viruses, as BtKY72 exhibited the weakest entry with *R. sinicus (472)* and *R. pearsonii* ACE2s. BtKY72 could utilize the ACE2 from *R. ferrumequinum* while the clade 1 RBDs could not. Khosta-1 exhibited compatibility with *R. ferrumequinum* ACE2, consistent with its identification in *R. ferrumequinum* bats[41]. The remaining clade 3 RBDs showed limited ACE2 compatibility aside from the sequence from *R. alcyone,* which permitted infection by all clade 3 virus RBDs tested.

### Interpretations of clade 3 *sarbecovirus* host range and tropism

Despite the various clade 3 sarbecovirus spike RBD and *Rhinolophus* ACE2 ortholog compatibilities observed in our pseudovirus infection assay, an important practical consideration is whether the virus is likely to encounter the corresponding bat in real life. For example, RhGB01, BB9904, and Khosta-2 were isolated from *R. hipposideros* or *R. euryale* bats in Europe (**Fig 3B, bottom**). Neither of these bats have been observed in Sub-Saharan Africa, and thus unlikely to allow transmission of these viruses to *R. alcyone* bats despite their potential RBD-ACE2 molecular compatibility. In contrast, BM48-31 was found in *R. blasii* bats, which have been observed in Eastern and Southern Africa, and thus in closer proximity to the known range of *R. alcyone* bats (**Fig 6A, top**). Thus, while all four RBDs can use the ACE2 from *R. alcyone,* only BM48-31 was found within a host bat of potentially overlapping geographical range (**Fig 6A, bottom**), and more likely to allow infection in those bats in real life.

**Figure 6.**
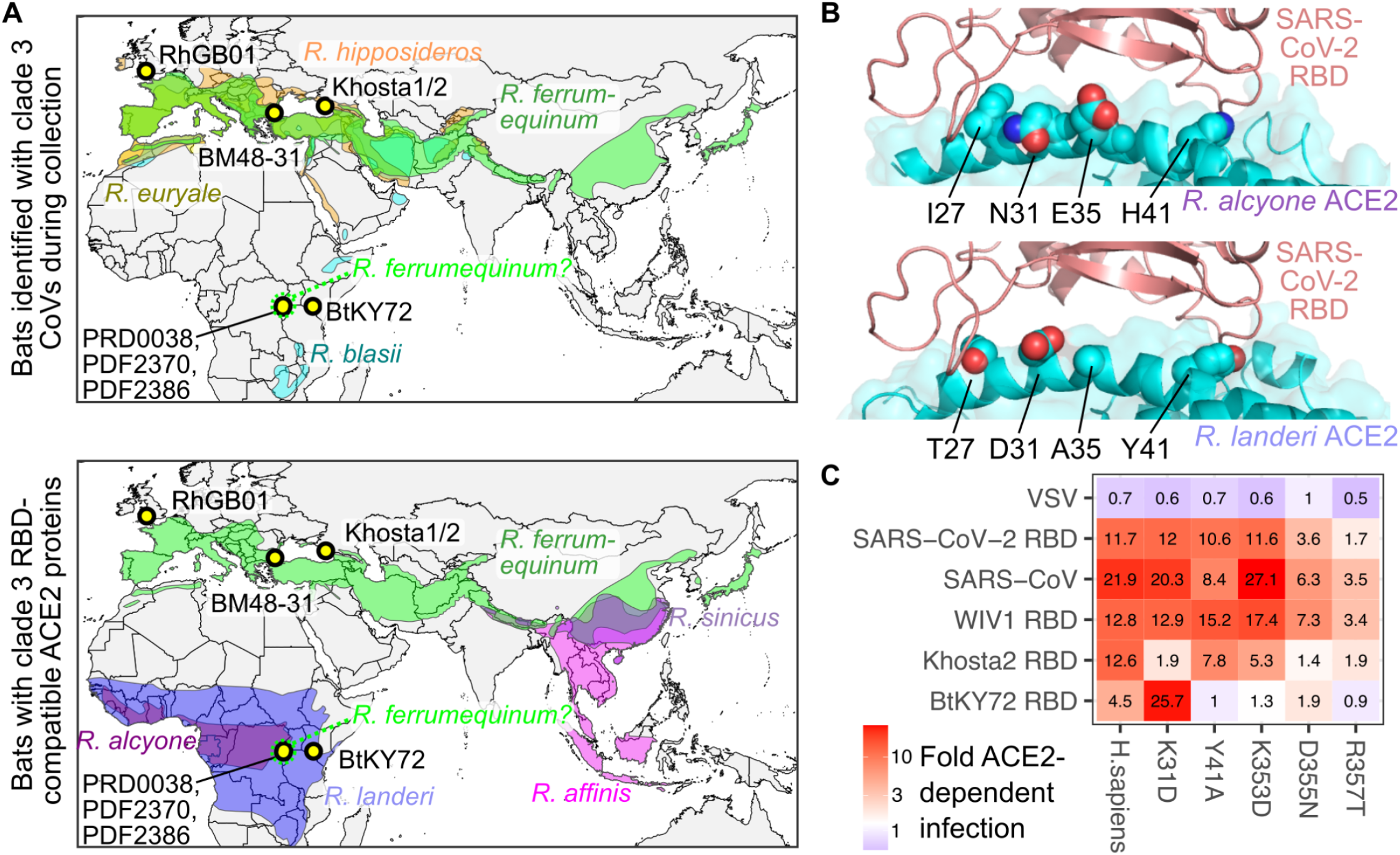
Rhinolophus bat ranges, collection sites, and interface residues. A) Map of known ranges of *Rhinotophus* bat species identified to harbor clade 3 sarbecoviruses at the time of sample collection (top), and bat species with potentially compatible ACE2 sequences empirically determined through pseudovirus infection assays (bottom). The geographic coordinates of sites of clade 3 sarbecovirus collections are shown as labeled yellow points with thick black outlines. B) The four RBD-interface residues differing between *R. alcyone* (top) and *R. landed* (bottom) ACE2 orthologs shown from RBD models based on PDB 6m17. C) ACE2-dependent infection measured by flow cytometry observed with RBD chimeric spike pseudoviruses in cells overexpressing mutants of human ACE2. All constructs encode human TMPRSS2 C-terminally linked to ACE2 with a 2A translational stop-start sequence.

Unlike the viruses isolated in Europe, BtKY72 and the related PDF-2370, PDF-2386, and PRD-0038 viruses were observed in *Rhinolophus* bats in Central and East Africa (**Fig 6A**). The exact species was not determined upon collection, although sequencing of the ACE2 gene from the PDF-2370 sample revealed that the host bat was likely *R. ferrumequinum* or a highly related species, only differing by 4 to 10 amino acids depending on the *R. ferrumequinum* ACE2 allele (**SFig 3**). *R. ferrumequinum* bats are not thought to populate Africa, although the host bat ACE2 sequenced from the PDF-2370 sample would suggest that *R. ferrumequinum,* or a highly related bat, is present in Central Africa. The next most similar of the currently sequenced ACE2 orthologs belonged to *R. alcyone* and *R. landeri,* differing by 25 and 28 residues, respectively (**SFig 3**). ACE2 sequences from *R. ferrumequinum, R. alcyone,* and *R. landeri* were all permissive for entry by BtKY72 RBD in our pseudovirus infection assay (**Fig 5**). The sites of sampling for all four viruses overlapped with the known range of *R. landeri* bats, and the more western sites where PDF-2370, PDF-2386, and PRD-0038 were sampled were at the edge of the known range of *R. alcyone* bats (**Fig 6A, bottom**). Thus, the BtKY72 related viruses are likely able to enter cells of various bat species, including *R. ferrumequinum* bats, around the sites they were first identified.

We also looked for ACE2 sequence patterns consistent with a history of sarbecovirus infection in these African horseshoe bats. The ACE2 proteins from *R. alcyone* and *R. landeri* are similar, differing only in 10 of the 805 total residues. Three of these differences are in the transmembrane domain and cytoplasmic tail, so our tested constructs only differed at seven positions. Four of these differences were at positions 27, 31, 35, and 41, all on the same surface of the main ACE2 alpha-helix that contacts the CoV RBDs (**Fig 6B**). While BtKY72 and Khosta-1 can use ACE2 proteins from both *R. alcyone* and *R. landeri,* Khosta-2, BM48-31, and RhGB01 are only capable of using the sequence from *R. alcyone* (**Figs 5A and 5C**), suggesting that these coding differences are functionally important for restricting entry for a subset of clade 3 sarbecoviruses. No sarbecoviruses have been identified in *R. alcyone* or *R. landeri* bats so far, but these genomic signatures suggest that these bats have been under evolutionary pressure by ACE2-utilizing sarbecoviruses in Africa.

### ACE2 lysine 31 impacts a subset of clade 3 RBD infections

Better understanding of the ways in which diverse sarbecoviruses have achieved compatibility with human ACE2 may help identify molecular barriers initially posed by ACE2 sequence differences that can be circumvented through RBD sequence adaptations from prior infection in various hosts. Within our set, the four RBDs capable of binding human ACE2 exhibited different patterns in *Rhinolophus* ACE2 ortholog usage. While they all recognize the same overall binding site on ACE2, they are likely presenting unique interfaces for the interaction, and thus relying on a different set of side chains within the same human ACE2 protein sequence. Our panel of *Rhinolophus* ACE2 orthologs differ at many residues, so it is impossible to delineate which specific amino acids are playing the most important roles in each interaction.

To gain amino-acid level resolution into this interaction, we tested a panel of human ACE2 mutants. We originally tested constructs that yield low amounts of overall ACE2 protein, as we previously saw that high ACE2 levels can mask the impacts of human ACE2 mutants during SARS-CoV or SARS-CoV-2 entry. These RBDs use the human ACE2 protein surface differently, as human ACE2 mutants Y41A and E37K reduced SARS-CoV spike-mediated entry without impacting SARS-CoV-2 [12]. We repeated the original singleplex experiments with the duplex infection assay (**SFig 7A**) and saw a strong correlation between the two assay results (Pearson’s r: 0.95, n = 30; **SFig 7B**).

Testing the same panel of mutants with the BtKY72 RBD was uninformative (**SFig 7A**), as this interaction is likely relatively weak and requires higher ACE2 amounts to promote infection. We thus recreated a smaller set of human ACE2 mutants in our high abundance ACE2 and TMPRSS2 co-expression construct, recombined them into HEK 293T landing pad cells, and exposed the cells to pseudoviruses with these RBDs (**Fig 6C, SFig 7C, D**). Both flow cytometry and microscopy readouts correlated well (**SFig 7E**). Most mutants had little impact on pseudovirus infection with SARS-CoV and SARS-CoV-2 RBDs in this context, except for D355N and R357T, which measurably reduced infection by all three clade 1 RBDs (**Fig 6C**), consistent with previous results[12].

The clade 3 sarbecoviruses had distinct reliances on human ACE2 mutants, typified by differences in how they were impacted by the K31D mutant (**Fig 6C**). Infection by both clade 3 RBDs were hampered by the D355N and R357T mutants, showing that these contacts serve a common role across diverse RBD interfaces. The Khosta-2 RBD was unaffected by the Y41A and K353D mutants, but particularly reliant on K31, as the K31D mutant reduced infection as strongly as D355N and R357T. In contrast, BtKY72 was strongly inhibited by most of the human ACE2 mutants tested including Y41A and K353D, with the sole exception being K31D, which drastically enhanced pseudovirus infection with the BtKY72 RBD (**Fig 6C**).

Despite the known signatures of evolutionary positive selection driving variation at ACE2 position 31 within bats (**Fig 4A**) [43], it is unclear which types of RBDs may be involved in this coevolutionary interplay. Clade 1 sarbecoviruses including SARS-CoV and SARS-CoV-2 are inhibited by the K31D mutant when ACE2 abundance is limiting (**SFig 7A**), suggesting that they either favor a Lys at this site or disfavor an Asp. Multiple co-structures show that SARS-CoV-2 spike Q493 can form a hydrogen bond with K31 in human or pangolin ACE2 (**SFig 8**) [31, 51], although this was not observed in all structures.

**Figure 7.**
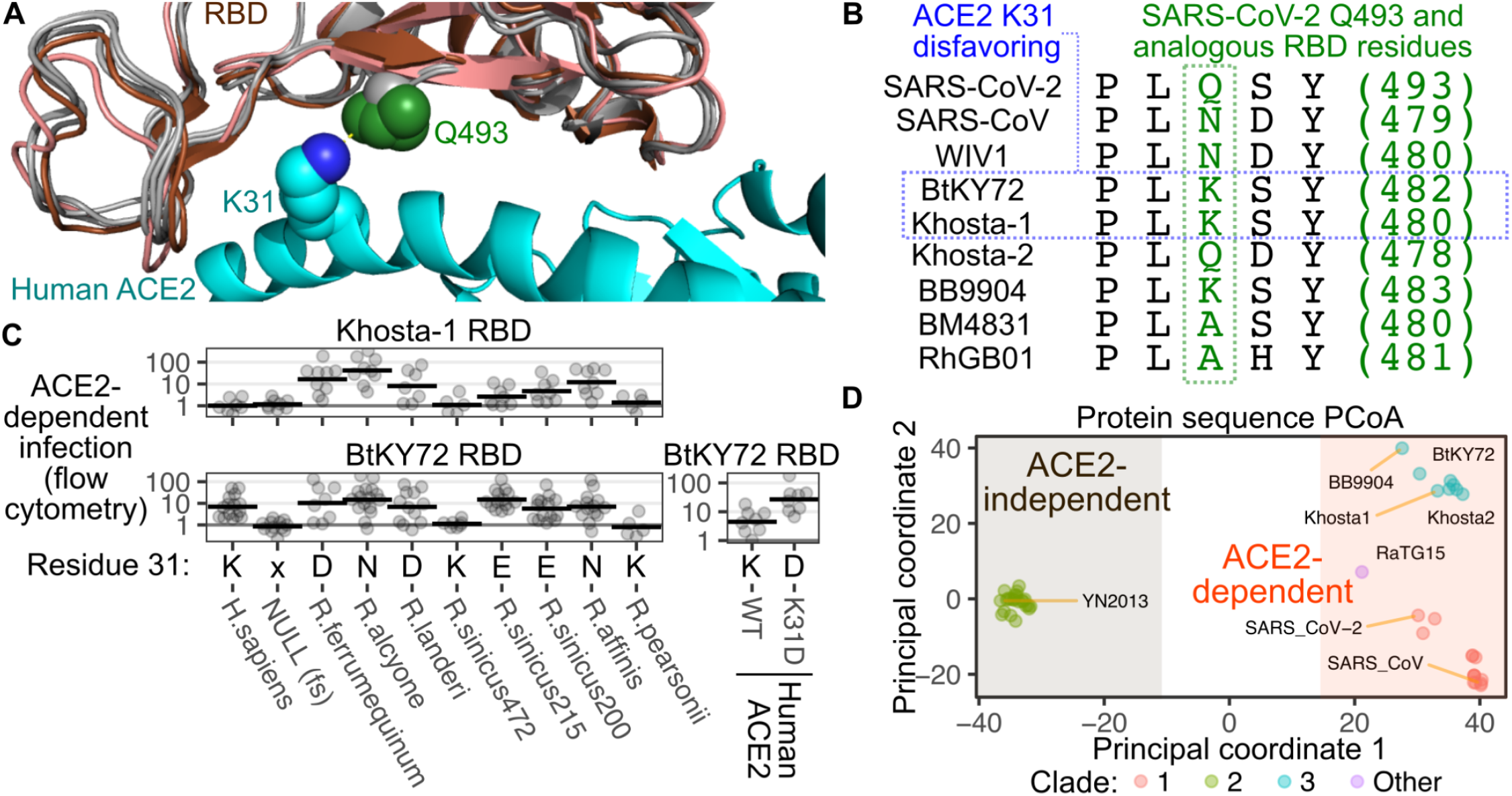
Impacts of ACE2 residue 31 on BtKY72 and Khosta-1 binding. A) The SARS-CoV-2 RBD (salmon) and human ACE2 interface (cyan; pdb: 6m17), overlaid with the SARS-CoV structure (brown, pdb: 7eam), and the BtKY72, Khosta-1, or Khosta-2 RBDs homology models (each shown in grey). The K31 residue on human ACE2 is shown as cyan spheres with the nitrogen group colored blue. The gamma-carbon atom of the RBD residue contacting ACE2 residue 31 is shown as a green sphere. B) Alignments of RBD residues including and adjacent to SARS-CoV-2 spike Q493, which extends from a conserved beta-strand and interacts with position 31 on ACE2 as shown in panel A. C) Comparison of patterns in ACE2 ortholog usage for BtKY72 and the side-chain present in ACE2 position 31. Human TMPRSS2 was also stably expressed in these cells. D) The Hamming distance matrix for sarbecovirus RBD protein sequences was used for a principal coordinate analysis. The first two principal coordinates are shown as X and Y axes, generating a scatter plot based on RBD sequence dissimilarity. Proposed delineation of ACE2-dependent and -independent RBDs based on differences in protein sequence are shown, with each clade given a different color.

To gain better insight at this molecular interaction, we aligned the homology models for BtKY72, Khosta-1, and Khosta-2 with the SARS-CoV and SARS-CoV-2 RBDs. As the SARS-CoV-2 Q493 position is at the end of a highly conserved beta-strand within the receptor binding motif, the models predicted the different amino acid side-chains at this position to extend from the same site and overall direction (**Fig 7A**). WIV1 and Khosta-2, which are both inhibited by the K31D human ACE2 mutant, encode either a Asn or Gln at this position and thus may also form similar hydrogen bonds as observed with SARS-CoV-2 (**Fig 7B**). BtKY72, which prefers the K31D human ACE2 mutant, encodes a Lys at this position (**Fig 7B**). This Lys side chain would lack the ability to hydrogen bond with K31 and their like charges may make close contacts energetically unfavorable.

We hypothesized that the pairwise identities of ACE2 residue 31 and the nearby residue on the RBD may determine compatibility for a subset of interactions. We tested this observation by looking at the ACE2 ortholog compatibility for BtKY72 and Khosta-1 (**Fig 7C**). Both of these viruses could not enter cells expressing Rhinolophid ACE2 orthologs encoding Lys at this position, while they could enter orthologs encoding Asp, Glu, or Asn. Khosta-1 was incapable of using human ACE2, which also encodes Lys at this position. The BtKY72 RBD could use human ACE2 thus defying this pattern, although its enhancement with the K31D mutant of human ACE2 supports our overall hypothesis (**Fig 7C**). The only other RBD with a Lys at this position was from BB9904. While this RBD was unable to support entry into cells expressing the K31 ACE2 orthologs from humans, *R. sinicus(472)* bats and *R. pearsonii* bats, this RBD also could not enter cells with other orthologs, and was less conclusive.

Altogether, our studies show that currently known sarbecoviruses have segregated into two groups of RBDs based on their ACE2-dependence or independence, with the more diverse ACE2-dependent viruses further segregating into sequence subgroups that each differentially utilize host ACE2 protein sequences during viral entry (**Fig 7D**). The sequences within the ACE2-dependent RBDs can be highly divergent, but a constellation of pairwise interactions, such as those between ACE2 position 31 and the adjacent RBD residue often encoding Gln, Asn, Lys, or Ala, likely determine the patterns of ortholog-specific compatibilities that enable successful entry during potential zoonotic events.

## Discussion

Here, we created a duplex pseudovirus infection assay for interrogating protein sequences capable of promoting viral infection. We subsequently harnessed this assay to test a matrix of 108 pairwise combinations for pseudovirus infection, with 12 different spike RBD sequences and 9 different ACE2 orthologs, to demonstrate that clade 3 sarbecoviruses consistently use various subsets of ACE2 alleles from a panel of horseshoe bats, with at least two clade 3 sarbecovirus spike RBDs also capable of using human ACE2. Our results also provide context for the importance of ACE2 residue 31, known to exhibit strong signatures of positive selection across bat species.

Our work was aided by the internally controlled, duplex nature of the infection assay format. This assay format recapitulated the effect size of the traditional singleplex format when the highly infectable cells were a minor fraction of the overall mixed cell population, and the magnitude of the effect was reduced by one-third when the cells were mixed equally. Testing the control and experimental samples in the same well allowed us to reduce the total number of samples in each experiment by up to two-fold. Notably, *in vivo* infections involve complex cell mixtures, and while the duplex infection assay does not fully recreate such conditions, it may be an improved proxy over traditional singleplex measurements for obtaining infection measurements. Future advancements with multiplexed assessments of receptor sequences in pseudovirus infection assays will likely come through uniquely barcoding each transgenic construct, so that cells that are sensitive or resistant to infection can be sorted and subsequently counted using high-throughput sequencing.

Viral glycoproteins that facilitate entry at the cell surface can cause cell-cell fusion resulting in syncytia formation. Consistent with other studies[52], we saw rampant syncytia formation with many of the WT or chimeric sarbecovirus spike proteins we tested, particularly when ACE2 and TMPRSS2 were both overexpressed. This observation prompted us to develop an automated microscopy readout for the duplex infection assay, as this does not require disrupting syncytia prior to measurement. While the dynamic range of our microscopy readout was smaller than the flow cytometry assay, the overall patterns were highly correlated between measurement types across all experiments. Our fluorescent nuclei markers aided the image-based analysis pipeline, as nuclei are far more consistent in size and shape than the cell bodies of adherent cells.

Despite the additional proteins proposed to serve as alternative receptors for SARS-CoV-2 infection, our side-by-side comparison showed ACE2 to confer the vast majority of enhancement to infection, followed by L-SIGN. While we did not test DCSIGN or SIGLEC1 [18, 53], these factors will likely confer similar effects as L-SIGN as they likely share a common mechanism for enhancing viral attachment to target cells through glycan binding. We did not see enhanced entry from overexpression of CD147, NRP1, or NRP2. Multiple recent studies have also observed no effect through CD147 overexpression [16, 17, 53], casting major doubt on its importance during SARS-CoV-2 entry and infection. Similar to our work, another study also did not observe significant effects from NRP1 or NRP2 overexpression[53], so the roles of these proteins during SARS-CoV-2 infection remain unclear.

Our sequence-function analysis revealed clear protein sequence and feature differences separating the ACE2-dependent and ACE2-independent groups. The ACE2-independent group completely overlaps with the evolutionary defined clade 2 sarbecoviruses. These RBDs are highly related to each other, but clearly distinct from the ACE2-dependent RBDs in amino acid length and sequence identity. In contrast to the structurally resolved clade 1 RBDs, including those from SARS-CoV and SARS-CoV-2, the ACE2-independent RBDs are ~ 15 residues shorter and are predicted to lack the disulfide bridged receptor binding ridge, particularly as the majority of ACE2-independent RBDs do not encode cysteines in this region of the interface. None of these RBDs conferred ACE2-dependent infection, even with sequences derived from bats of the same species as the ones they were isolated from.

In contrast, the ACE2-dependent RBDs included all known clade 1 and clade 3 viruses, and the viruses within the recently discovered RaTG15 clade[30]. These RBDs were between 219 and 223 residues in length, possessed a pair of cysteines in the receptor binding ridge capable of forming a disulfide bridge, and exhibited species-specific utilization of at least one known rhinolophid ACE2 protein. Despite their shared feature of ACE2 utilization, these RBDs can still drastically vary in protein sequence, with diverse pairs of RBDs exhibiting amino acid differences at 50 to 75 positions (**SFig 1**). While all known clade 1 and clade 3 sarbecovirus RBDs share these features, there are undoubtedly additional clades of sarbecovirus RBDs not yet observed which may defy these patterns.

Two other preprints have recently observed similar findings with clade 3 sarbecovirus RBD utilization of ACE2. Starr and colleagues observed weak *in vitro* binding between BtKY72 RBD and human ACE2 [54], which could be enhanced with a K493Y/T498W double mutant in the RBD that increases its affinity. They, too, were initially unable to observe pseudovirus infection with the full length WT BtKY72 spike, although the double mutant spike yielded detectable infection [54]. Seifert and Letko observed that the Khosta-2 RBD could use human ACE2 during infection [55]. Both of these observations are consistent with our results, considering slight differences in assay sensitivities. By generating a larger set of rhinolophid ACE2 orthologs including *R. ferrumequinum, R. alcyone,* and *R. landeri,* we were also able to test sequences that were likely more similar to those in the bats that serve as their natural hosts, instrumental in seeing that all of the known sarbecoviruses with RBDs possessing 219 or more residues are ACE2-dependent. The precise genetic determinants of compatibility between spike RBD and host ACE2 sequences will become clearer once more *Rhinolophus* ACE2 sequences are sequenced and tested in functional assays like ours.

Despite the overall concordance in results from independent groups, some inconsistencies remain. For example, a previous study by Wells and colleagues concluded that the PRD-0038 and PDF-2370 / PDF-2386 RBDs, which they refer to as “Rwanda” and “Uganda” viruses, do not utilize human ACE2 [11]. These RBDs are highly similar to the clear human ACE2-compatible BtKY72 RBD, differing only in 3 or 4 amino acids across the ~ 222 residue RBD. It is currently unclear whether this discrepancy is due to a difference in assay sensitivity, or whether the relatively small number of amino acid differences between these highly related RBDs were capable of drastically altering their affinity for human ACE2.

Even with the advantages of our assay and results, interpretations should be made with caution. Due to financial constraints, the diverse sarbecovirus spikes and *Rhinolophus* ACE2 alleles we tested were chimeric molecules, wherein the domains most critical for the virus-host interaction were swapped into existing scaffold protein sequences. As some chimeric molecules may not be fully stable, there are the possibilities of false negatives in our dataset. For example, the strongest enhancement to pseudovirus infection conferred by *R. sinicus* (472) ACE2 cells was the 3.5-fold increase with WIV1 chimeric spike, and it is currently unclear whether this relatively poor enhancement is a true property of this allele, or an artifact of altered protein conformation or subcellular localization.

While BtKY72 and Khosta-2 RBDs can utilize human ACE2 during entry, this does not mean that these viruses are currently capable of infecting humans. In both cases, the amounts of pseudovirus infection conferred by these RBDs were less than those conferred by the SARS-CoV, SARS-CoV-2, and WIV1 RBDs. These interactions may still be too inefficient to allow wide-spread entry and replication within humans. Furthermore, replication *In vivo* is multifactorial[56], and there are likely additional incompatibilities in immune antagonism and replication that may stifle a zoonotic event. For example, the Khosta viruses lack genes thought to antagonize the immune system, such as ORF8[41]. While not sufficient, compatible interactions between viral entry proteins and host receptor proteins are likely necessary for zoonosis. Thus, these results demonstrate that there are sarbecoviruses that are at least partially primed to jump into humans, and that surveillance efforts should be further extended outside of East Asia to other continents, including Africa and Eastern Europe.

The drastic differences in ACE2 and sarbecovirus RBD compatibility observed in our study highlight the importance of knowing the genotypes of both the virus and host. For example, *R. alcyone* was compatible with all clade 1 and 3 RBDs tested, while the highly related ortholog from *R. landeri* was only compatible with half of these viruses, and the next related sequence from *R. ferrumequinum* was only compatible with BtKY72 and Khosta-1. All three species may overlap in geographical range in Central Africa, and accurate identification of the host bat is needed to know which species can serve as reservoirs *In vivo.* There is a staggering amount of ACE2 allelic variation in horseshoe bats, including the 19 or more variants observed with *R. sinicus,* the 6 or more variants observed with *R. affinis*, and 3 or more variants observed with *R. ferrumequinum.* With some pairs of alleles in the same species differing by as much as 17 residues, there are likely drastically different viral susceptibilities and phenotypic heterogeneity within a population of the same species. Future efforts capable of simultaneously sequencing both the viral genome and host ACE2 coding sequence from a single sample will undoubtedly help uncover some of these complex relationships between virus and host.

Our results clarify the molecular interactions that likely underlie the evolutionary interplays between sarbecovirus RBDs and host ACE2 sequences. For example, ACE2 residue 31 was long known to be a site of evolutionary conflict due to its signature of positive selection in bats[43]. We found that the BtKY72 and Khosta-1 RBDs disfavor horseshoe bat ACE2 orthologs and alleles encoding K31. Human ACE2 also encodes K31. While BtKY72 was capable of using human ACE2, its infection was enhanced with K31D mutant human ACE2. Both RBDs encode Lys at the RBD residue that interacts with ACE2 position 31, suggesting a potential side-chain incompatibility making the interaction less energetically favorable.

Consistent with our interpretation, Starr and colleagues found that the WT BtKY72 Lys residue was disfavored for human ACE2 interaction, as substitution to multiple other amino acids including Tyr, Gln, Phe, Ala, Val, Gly, and Cys improved binding[54]. In contrast, SARS-CoV and SARS-CoV-2 RBDs encode Asn or Gln at the analogous residue, and disfavor the K31 D mutant of human ACE2. Thus, an amino acid that is favored for one sarbecovirus RBD is disfavored for the other, and *vice versa*. A similar pattern of incompatibility was previously observed between N479 of SARS-CoV RBD or K479 from a related virus cSz02 from palm civets, and K31 of human or T31 of palm civet ACE2, thought to be important for transmission of SARS-CoV from palm civet intermediate hosts[6]. Similar interplays likely exist between other key pairs of RBDs and ACE2 residues, and the collective actions of all of these interactions likely dictate ACE2 usage by a given RBD.

The current pandemic is a sobering reminder of the importance of understanding the molecular barriers that normally prevent zoonosis, especially when previous ecological and societal factors that also served as barriers continue to erode. Once identified, weakened barriers can be bolstered or surveilled as part of pandemic precaution. While most prescient with sarbecoviruses, these considerations apply to other viruses, including Merbecoviruses, Henipaviruses, or Filoviruses. Multiplexable genetic assays will be instrumental in getting the large number of data points needed to understand the nuanced molecular coevolutionary relationships that exist across a diverse set of host-pathogen interactions.

## Materials and Methods

### Plasmid Construction

Construction of the landing pad lentiviral vector construct, LLP-Int-BFP-IRES-iCasp9-Blast (Addgene plasmid #171588) was described previously[12]. All plasmids were produced using Gibson Assembly[57]. For the initial polymerase chain reaction, a total of 40 ng template plasmid DNA was mixed with forward and reverse primers, each at a final concentration of 0.333 μM, and amplified with Kapa HiFi HotStart ReadyMix polymerase. To create the chimeric SARS-CoV spike constructs with chimeric RBDs, 100 ng of gBlock DNA (Integrated DNA Technologies) encoding the codon-optimized RBD sequences were used as the starting template. To create the chimeric rhinolophid ACE2 molecules, 100 ng of eBlock DNA oligomers (Integrated DNA Technologies) were used as the template. Notably, the multi-piece eBlock ligation strategy worked poorly, and is not recommended for future molecular cloning.

All of the aforementioned DNA was amplified under the following conditions: 95C’ 5’, 98C’ 20”, 65C’ 15”, 72°C 8’, repeat seven or eight times, 72°C 5’. Twenty units (1 μL) of DPN1 enzyme (New England BioLabs, R0176L) were added to each reaction, except for those produced from DNA oligomers, and incubated for two hours at 37°C. A Zymo clean and concentrator kit (Zymo Research, D4003) was used to clean each reaction and 1 μL of the final eluate was incubated with 1 μL 2x GeneArt™ Gibson Assembly MasterMix (ThermoFisher, A46629) for 30-60 minutes at 50°C to complete the Gibson cloning reaction. The resulting recombinant plasmids were transformed via calcium heat shock into home-made chemically-competent E. coli 10β cells (New England BioLabs, C3019I). Plasmid DNA was extracted using a GeneJET miniprep kit (ThermoFisher, K0503) and sequence-confirmed with Sanger sequencing on an Applied Biosystems 3730 Genetic Analyzer.

The plasmid backbones for the aforementioned molecular cloning were described in our previous publication [12], including AttB_ACE2_IRES-mCherry-H2A-P2A-PuroR (Addgene plasmid #171594), AttB_[Kozak-mut]ACE2_IRES-mCherry-H2A-P2A-PuroR (Addgene plasmid #171595), and AttB_ACE2[dEcto]_IRES-mCherry-H2A-P2A-PuroR (Addgene plasmid #171596). psPAX2 and pMD2.G were gifts from Didier Trono, (Addgene plasmid #12260; http://n2t.net/addgene:12260; RRID: Addgene_12260) and (Addgene plasmid # 12259; http://n2t.net/addgene:12259; RRID:Addgene_12259), respectively. Plasmids encoding coronavirus spike proteins for MERS-CoV, SARS-CoV, SARS-CoV-2 (Wuhan-Hu-1) and WIV1-CoV were a kind gift from David Veesler. The plasmids encoding the viral glycoproteins for Ebolavirus Zaire, Marburgvirus, Lassa fever virus, VSV, Junin virus and LCMV were a gift from James Cunningham and have been previously described [27]. NRP1 coding sequence was amplified from MAC-NRP1 (Addgene plasmid #158384). The NRP2 coding sequence was amplified from DNASU plasmid HsCD00398541, CD209 was amplified from DNASU plasmid HsCD00779810, CLEC4M was amplified from DNASU plasmid HsCD00413491, and BSG was amplified from DNASU plasmid HsCD00849488. The RBD and ACE2 ortholog sequences used in our study are listed in **Supplementary Table 1**. R. sinicus (200) corresponds to NCBI accession QMQ39200.1, R. sinicus (215) corresponds to accession QMQ39215.1, and R. sinicus (472) corresponds to ADN93472.1.

### Cell culture

Cell culture reagents were purchased from ThermoFisher unless otherwise noted. Cell lines were cultured in D10 medium: Dulbecco’s modified Eagle’s medium supplemented with 10% fetal bovine serum (Gibco,10437028), 100 U/mL penicillin, and 0.1 mg/mL streptomycin (Corning, 30-002-CI). Cells were passaged via detachment with Trypsin-Ethylenediaminetetraacetic acid 0.25% (Corning, 25-053-CI). Landing pad cells were grown in D10 supplemented with 2 μg/mL doxycycline (Fisher, AAJ67043AD), indicated as D10-dox. Long-term passaging of landing pad cells utilized D10-dox with 20 μg/mL blasticidin (InvivoGen, ANT-BL-1) to remove cells with silenced landing pad loci.

### Recombination of landing pad cells

HEK 293T cells were used to generate the lenti-landing pad line derived from LLP-Int-BFP-IRES-iCasp9-Blast as previously described[10]. Landing pad cells expressing Bxb1 integrase with a nuclear localization signal to allow for transport into the nucleus were recombined in either 24-well or 6-well plates. In the 24-well plate, 120,000 cells were transfected with 254 ng of attB recombination plasmid mixed with 0.96 μL of Fugene 6 reagent in D10-dox media. In the 6-well plate, 600,000 cells were transfected with 1,200 ng of attB recombination plasmid mixed with 5 μL of Fugene 6 reagent in D10-dox media.

Upon attB-plasmid transfection, negative selection of non-recombined landing pad cells was performed with the addition of 10nM AP1903 (ApexBio, B4168) to activate iCasp9. Positive selection of recombined cells was achieved with the addition of 1 μg/mL puromycin (InvivoGen, ANTPR1). Recombined cells were maintained in D10-dox with 1 μg/mL puromycin to prevent transgene silencing.

### BLAST searches and protein sequence alignments

The receptor binding domains of SARS-CoV, WIV1, and SARS-CoV-2 spikes were used as initial query sequences for National Center for Biotechnology Information (NCBI) BLASTp searches. The resulting sarbecovirus spike protein sequences were obtained from NCBI, with the corresponding accession numbers listed in Supplementary Table 1. All RBD fragments were manually curated and aligned using Clustal Omega[58]. The spike RBD sequences for PDF-2370 and PDF-2386 were identical, and we thus collapsed these two entries into one and only refer to this sequence as PDF-2370 for simplicity. The aligned sequences were used as the input for a custom python script that performed calculations of amino acid identity at each position for any given pair of RBD sequences.

To perform a comprehensive search for clade 2 RBD spike sequences to gain a near-complete sampling of their sequence diversity, we first performed an NCBI BLASTp search using the YN2013 RBD amino acid sequence as the query. Clade 2 sequences were retained following a filtering step excluding hits longer than 210 amino acids, yielding a list of 112 likely “clade 2” accession numbers. The full “YN2013” spike amino acid sequence was then used to perform another search, and full length spike sequences were retrieved and separated based on their existence or absence in the list of “clade 2” RBDs.

To identify *Rhinolophus* bat ACE2 alleles, we performed an NCBI BLASTp search using human ACE2 as the query sequence but restricting results to the 58055 taxonomic ID. The sequences were aligned, and Hamming distance matrices were calculated, as described before for the sarbecovirus RBD sequences.

### Pseudotyped virus infection assays

All pseudotyped virus infection experiments were performed with lentiviral vectors. The lentiviral vectors were produced by transfecting 1.5 million HEK 293T cells in a single well of a 6-well plate, using PEI-Max MW 40,000 (PolySciences, CAS Number: 49553-93-7) mixed with 600 ng of PsPax2 (Addgene # 12260), 600 ng of the lentiviral transfer vector pLenti_CMV-EGFP-2A-mNeonGreen (Addgene # 171599), and 600 ng of various viral envelope plasmids. The media was changed the next day, and the supernatant was collected over the next 72 hours. Upon each collection, the media was spun at 300 × g for 3 min, and the soluble fraction retained. A list of viral envelope coding sequences used in this study is shown in Supplementary Table 1. Upon mixing of pseudovirus supernatants and target cells mixtures, the target cells were incubated for 48 or more hours prior to processing for flow, or imaging by automated microscopy.

For comparing the singleplex and duplex infection assays, 50,000 cells stably modified with the attB-ACE2(dEcto)_IRES-mCherry-H2A-2A-PuroR plasmid, 50,000 cells stably modified with the attB-ACE2_IRES-iRFP670-H2A-2A-PuroR plasmid, or various combinations of both cells were plated into individual wells of a 24-well plate. These plates were then mixed with ~ 10 uL of VSV-G pseudovirus, ~ 100 uL of SARS-CoV spike pseudovirus, and ~ 800 uL of SARS-CoV-2 spike pseudovirus. For the singleplex assay analysis, the percent of GFP positive cells in the sample modified with attB-ACE2_IRES-iRFP670-H2A-2A-PuroR was divided by the percent of GFP positive cells in the sample modified with attB-ACE2(dEcto)_IRES-mCherry-H2A-2A-PuroR, to get the fold ACE2-dependent infection. For the duplex assay, within the samples where there were mixtures of the two differentially modified cells, the percent of GFP positive cells within the iRFP670 positive population was divided by the percent of GFP positive cells within the mCherry positive population, to yield the fold ACE2-dependent infection from the cells within a single well. The proposed alternative receptor infection experiments were performed by plating ~ 50,000 cells total per well of a 24-well plate. For the experiments testing the matrix of RBDs and ACE2 alleles, 10,000 total cells were plated per well of a 96-well plate. To maximize viral titers, all viral supernatants were used fresh.

### Western Blotting

Pure populations of LLP-Int-BFP-IRES-iCasp9-Blast HEK 293T cells stably modified with the ACE2(dEcto) construct, HA-tagged proposed receptor constructs, or human or Rhinolophus ACE2 constructs were selected and maintained in D10-dox with 1 μg/mL puromycin. These cells were lysed using 1x RIPA buffer (Thermo Scientific, #89901) supplemented with 1X protease inhibitor (Thermo Scientific, #1862209). Protein lysates were quantified using BCA protein reagents A & B (Thermo Scientific, #23223, 23224) and 20 μg of protein lysates from each sample were denatured in 4X LDS sample buffer followed by separation on 4–12% gradient SDS PAGE polyacrylamide gel (Genscript, #M00653). Separated proteins on the polyacrylamide gel were transferred to a 0.2 μm PVDF membrane (Thermo Scientific, #88520) and immunoblotted using anti-ACE2 (Abcam, #Ab15348), anti-HA-HRP (3F10 antibody from Roche) or anti-β-actin antibody (Santa Cruz, #S47778). Western blot images were acquired using a GE Amersham Imager 600.

### Flow cytometry and fluorescence microscopy

Cells were detached with 0.25% Trypsin with 2.21 mM EDTA (Corning, #25-053-CI), and resuspended in PBS containing 5% fetal bovine serum. Analytical flow cytometry was performed either with a ThermoFisher Attune NxT or a BD LSRII flow cytometer. For the Attune NxT, mTagBFP2 was excited with a 405 nm laser, and emitted light was collected after passing through a 440/50 nm bandpass filter. EGFP was excited with a 488 nm laser, and emitted light was collected after passing through a 530/30 nm bandpass filter. mCherry was excited with a 561 nm laser, and emitted light was collected after passing through a 620/15 nm bandpass filter. iRFP670 and miRFP670 were excited with a 638 nm laser, and emitted light was collected after passing through a 720/30 nm bandpass filter. For the BD LSRII, mTagBFP2 was excited with a 405 nm laser, and emitted light was collected after passing through a 440/40 nm bandpass filter. EGFP was excited with a 488 nm laser, and emitted light was collected after passing through a B525/50 nm bandpass filter. mCherry was excited with a 561 nm laser, and emitted light was collected after passing through a 610/20 nm bandpass filter. iRFP670 and miRFP670 were excited with a 640 nm laser, and emitted light was collected after passing through a 710/40 nm bandpass filter. Before analysis of fluorescence, live, single cells were gated using FSC-A and SSC-A (for live cells) and FSC-A and FSC-H (for single cells).

Fluorescent images were captured on a Nikon Ti-2E fluorescent microscope, outfitted with a SOLA SM II 365 light engine (Lumencor), a CFI Plan Apochromat DM Lambda 20X objective or a NIKON Plan Fluor 4X objective, GFP (#96392), Texas Red (#96395), or Cy5 (#96396) filter sets, and imaged with a DS-QI2 monochrome CMOS camera. The images were captured with an automated image acquisition workflow, which performed autofocus on each well of a 96-well plate. All exposure times for each fluorescent channel were kept constant between wells and replicate experiments. The captured TIFF files were analyzed with a custom Python script utilizing the numpy, scipy, cv2, skimage, and PIL packages. This script entitled “Overlap_ratio_calculation.py” can be found in the project GitHub repository (https://github.com/MatreyekLab/ACE2_dependence). The image shown in Fig 5B was processed in NIS-Elements imaging software (Nikon), with intensity minimums and maximums auto scaled to show the locations and relative sizes of the red or near-infrared nuclei, and highlight the distribution of GFP within the large syncytial cell.

### Data Analysis and statistics

Data analysis was performed using version 1.4.1717 of RStudio, with the exception of flow cytometry data, which was first analyzed using version 10.8.0 of FlowJo. An R Markdown file containing code capable of fully reproducing the analyses can be found at the Matreyek Lab GitHub repository (https://github.com/MatreyekLab/ACE2_dependence). The analysis utilized the tidyverse[59], ggrepel, ggbeeswarm, sf[60], and ggfortify[61] packages. Statistical significance was determined using two-sided t-tests, and multiple test corrections were performed using the Benjamini-Hochberg procedure. The principal component analysis was performed by using the cmdscale classical (metric) multidimensional scaling function in ggfortify on the RBD Hamming distance matrix shown in SFig 1.

To calculate ACE2 dependent infection by flow cytometry, the acquired single cells were subsequently gated into mCherry+/iRFP670- or mCherry-/iRFP670+ sub-populations using FlowJo. The percentage of GFP positive cells in each subpopulation was calculated and exported as individual columns of a comma-separated value datafile. These values were copied into the experiment sample sheet listing the date, sample name, cell line used, pseudotyped virus used, and pseudotyped virus inoculum, and imported into RStudio for subsequent analysis. There the percent GFP value for the mCherry-/iRFP670+ subpopulation was divided by the percent GFP value for the mCherry+/iRFP670-population to obtain a ratio. The geometric mean of multiple replicate experiments were used to derive the ACE2-dependent infection metric used throughout our work.

### Modeling of Bat ACE2 three-dimensional structures

The HHpred web server was used to perform homology alignment of various Bat ACE2 sequences with human ACE2 structures (pdb: 6m17 and 6m18) [62, 63]. A structural model was then built with the MODELLER web server [64], and the ACE2 models were each aligned to ACE2 in PDB:6m17 using the default alignment settings in PyMol. The HHpred web server was used to perform homology alignment of the YN2013 RBD sequence with the SARS-CoV RBD (pdb: 7lm9). The HHpred web server was also used to align the RBD of the BtKY72 spike protein to the spike protein from SARS-CoV-2 (pdb: 7eam). A structural model was then built with the MODELLER web server [64], and the ACE2 models were each aligned to the SARS-CoV-2 RBD in PDB:6m17 using the default alignment settings in PyMol. The same pipeline was used to generate a model of the SARS-CoV-2 RBD, to gain an estimate of the amount of error produced in the modeling process.

### Visualizing the global ranges of various rhinolophus bat species

The ranges of R.affinis[65], R.alcyone[66], R.blasii[67], R.euryale[68], R. ferrumequinum[69], R.hipposideros[70], R.landeri[71], and R.sinicus[72] were downloaded as shape files from The IUCN Red List of Threatened Species 2020. The shape files were imported into R and displayed in R studio using the “sf” package[60].

## Acknowledgements

We wish to thank Simone Edelheit and Milena Zelembaba of the Genomics Core Facility of the CWRU School of Medicine’s Genetics and Genome Sciences Department.

## Author contributions

Conceptualization: Kenneth A. Matreyek.

Data curation: Kenneth A. Matreyek.

Formal analysis: Kenneth A. Matreyek.

Funding acquisition: Kenneth A. Matreyek, Anna M. Bruchez

Investigation: Sarah M. Roelle, Nidhi Shukla, Anh Pham, Anna M. Bruchez, Kenneth A. Matreyek.

Methodology: Kenneth A. Matreyek.

Project administration: Kenneth A. Matreyek.

Resources: Kenneth A. Matreyek.

Supervision: Kenneth A. Matreyek.

Visualization: Kenneth A. Matreyek.

Writing – original draft: Kenneth A. Matreyek.

Writing – review & editing: Kenneth A. Matreyek, Anna M. Bruchez, Nidhi Shukla, Sarah M. Roelle

## SUPPLEMENTAL FIGURES

**SFig 1.**
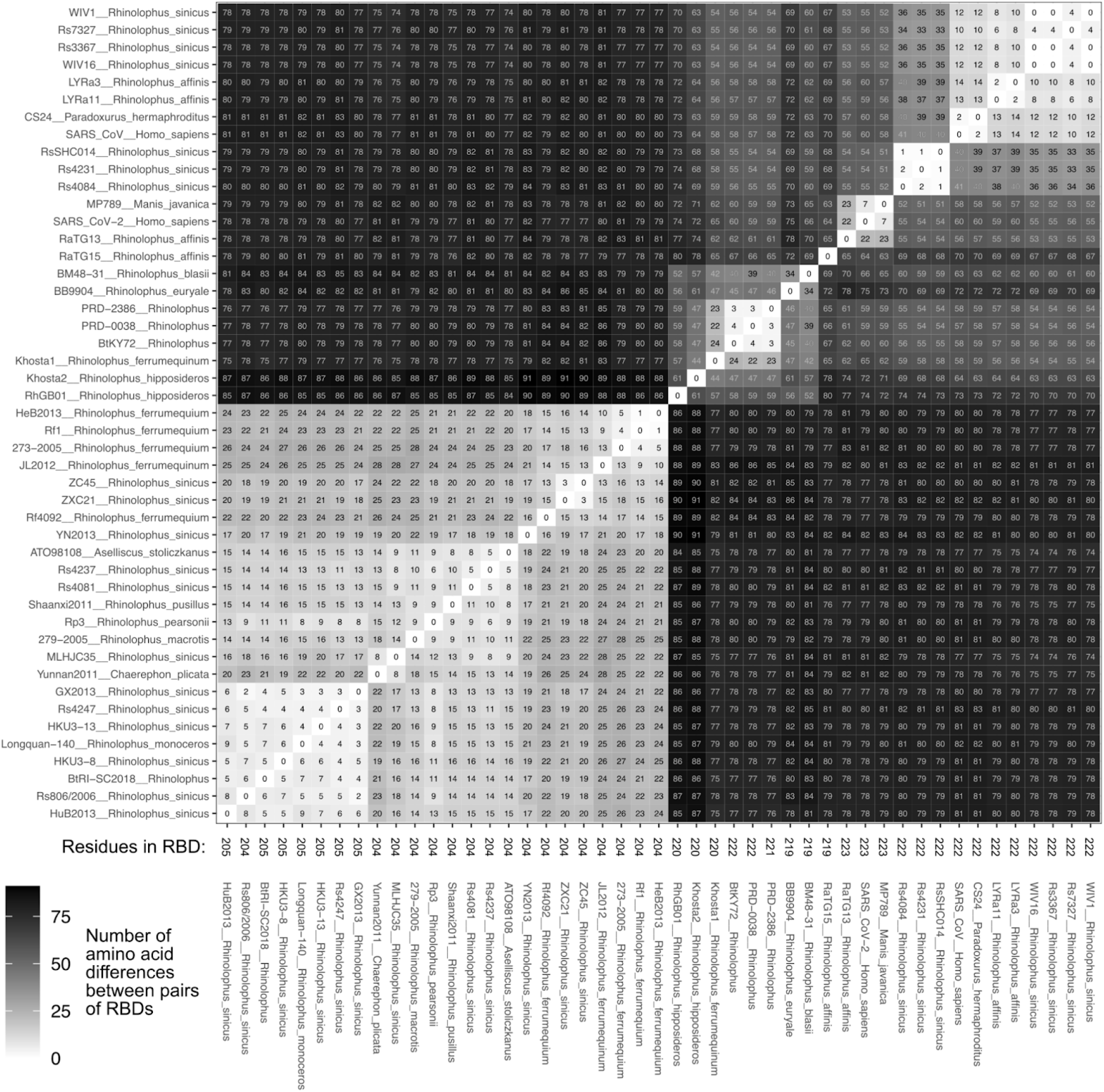
Hamming distance matrix of a representative diversity of sarbecovirus RBDs. Similar to Fig 3E, except a larger set of RBDs were used as input. While not all known RBDs are shown, the smallest subset of samples capturing the known diversity of RBD sequences were chosen. The numbers within the boxes denote the number of residues that differ between the pairs of sequences. The numbers along the bottom axis labels denote the total number of residues in the RBD.

**SFig 2.**
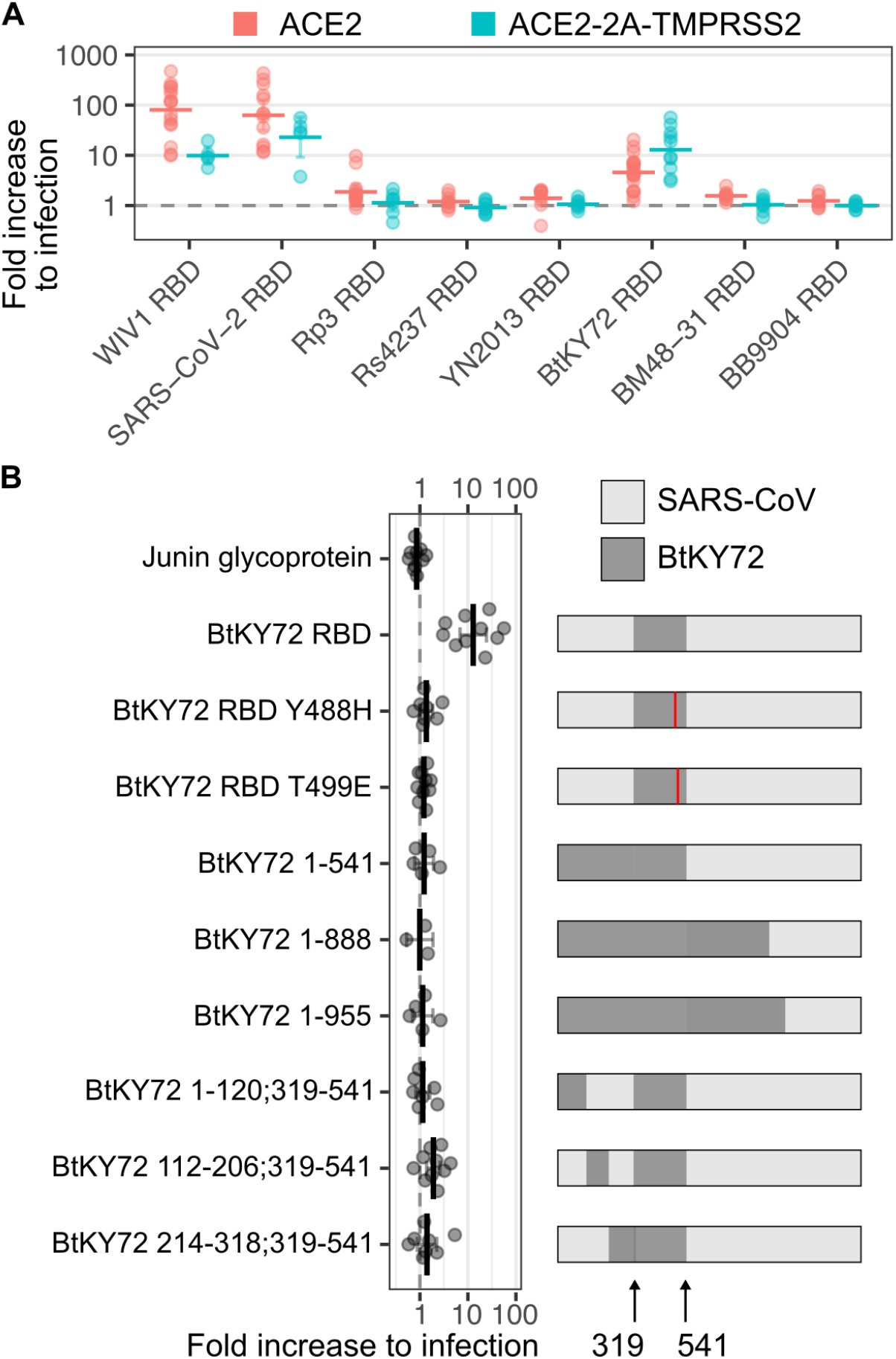
Additional figures of clade 3 RBD pseudovirus infectivities. A) Comparison of ACE2-dependent infectivities observed with HEK 293T cells overexpressing ACE2 only or ACE2 with TMPRSS2. B) Comparison of ACE2-dependent infectivities observed with various chimeric sequences made between BtKY72 and SARS-CoV spikes.

**SFig 3.**
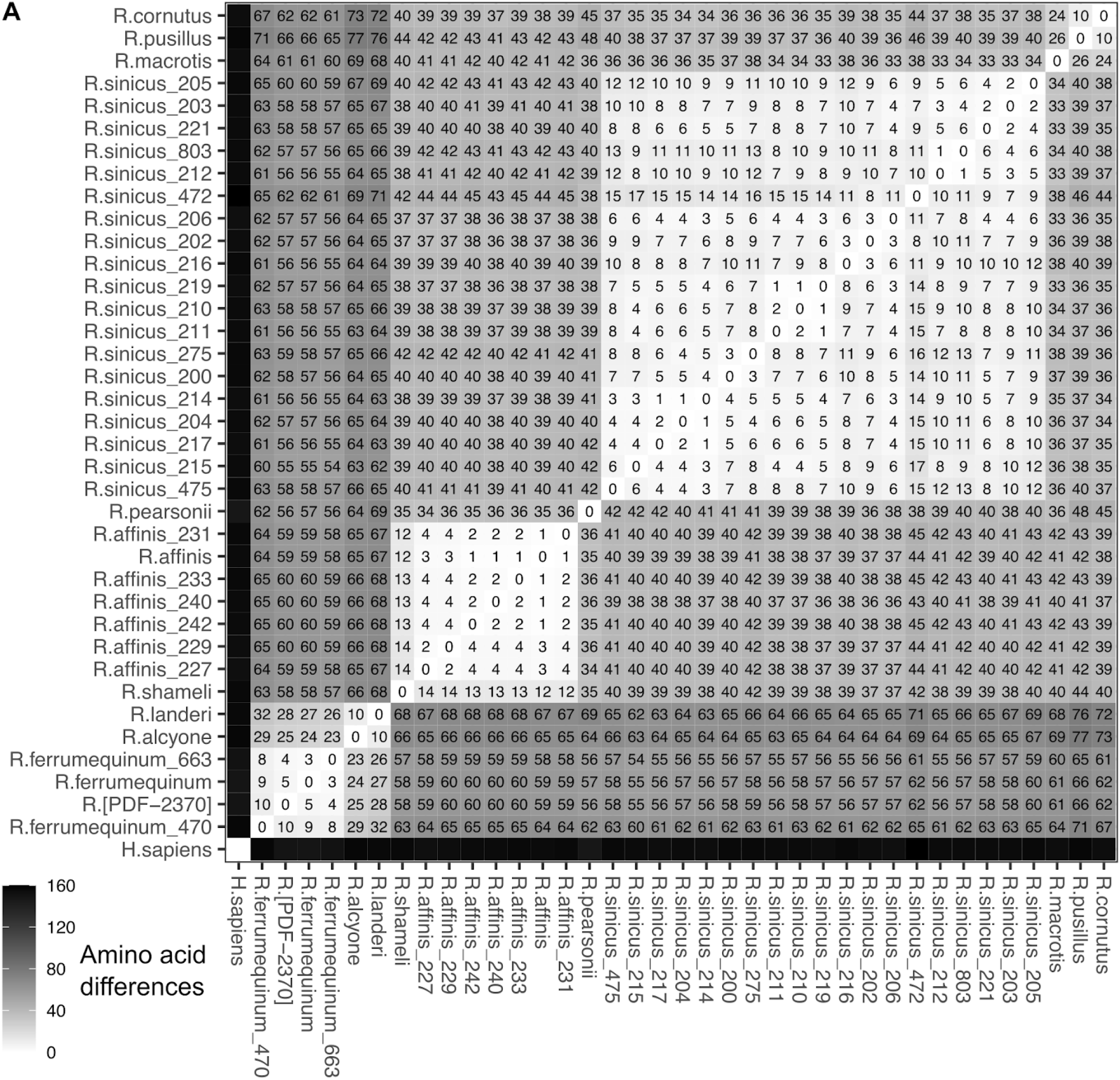
Hamming distance matrix of ACE2 sequences observed in various horseshoe bats. Unlike Fig 4C which shows the amino acid differences in the chimeric ACE2 proteins we tested, this figure shows the Hamming distances for the full length protein sequences.

**SFig 4.**
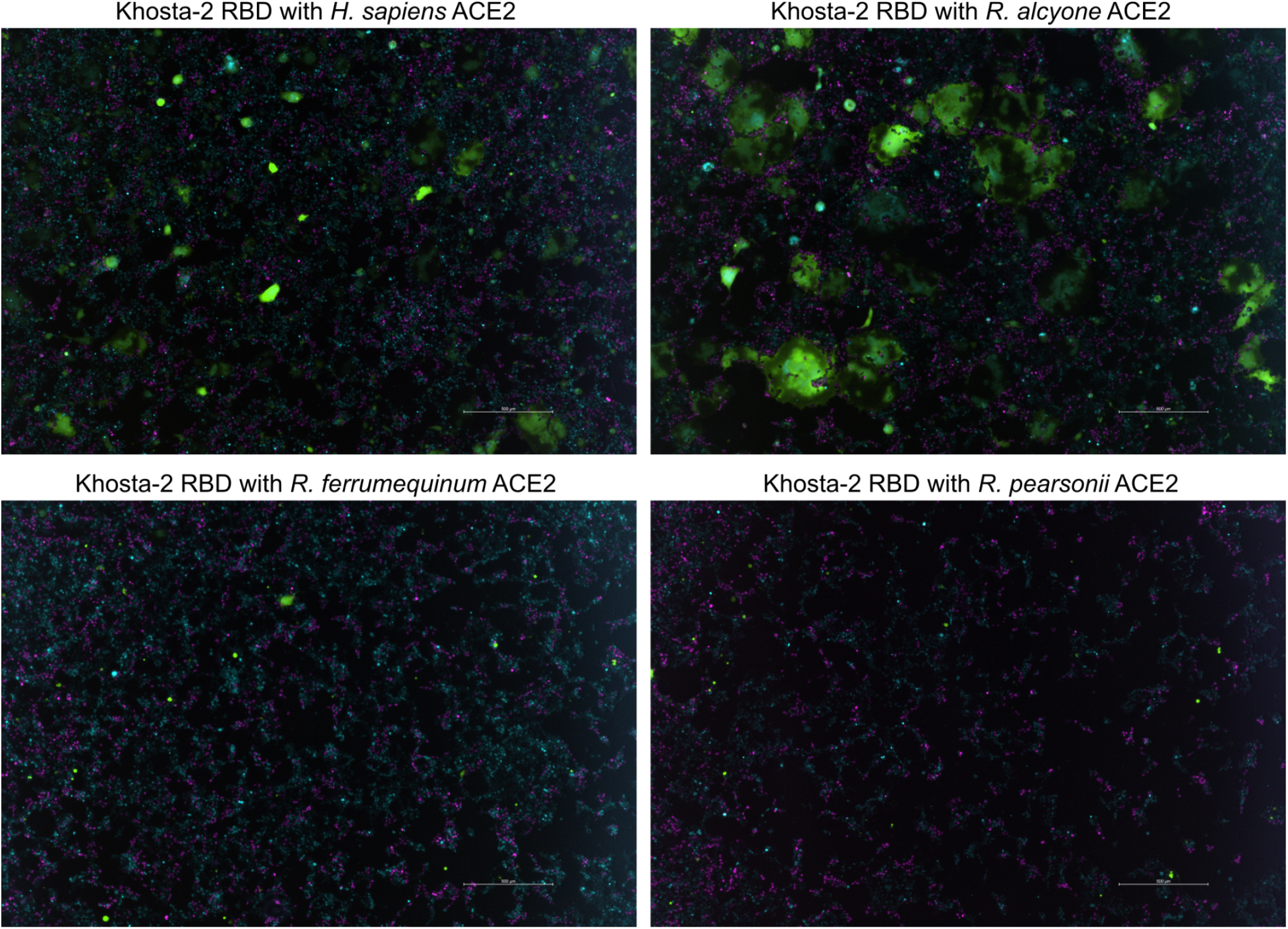
Images of syncytia when cells expressing various ACE2 orthologs were infected with Khosta-2 RBD chimeric spike pseudoviruses. Green fluorescence marks cell bodies from syncytia in which at least one cell had been infected by Khosta-2 RBD pseudovirus. Magenta dots are ACE2 negative control cell nuclei expressing mCherry-fused histone H2A, while cyan dots are ACE2 ortholog cell nuclei expressing iRFP670-fused histone H2A. Images were taken with a 4x objective, and with a 500 micron scale bar shown at the bottom right of the image. All experiments were performed with HEK 293T cells overexpressing the indicated ACE2 sequence and human TMPRSS2 co-translationally linked together with a 2A translational stop-start sequence.

**SFig 5.**
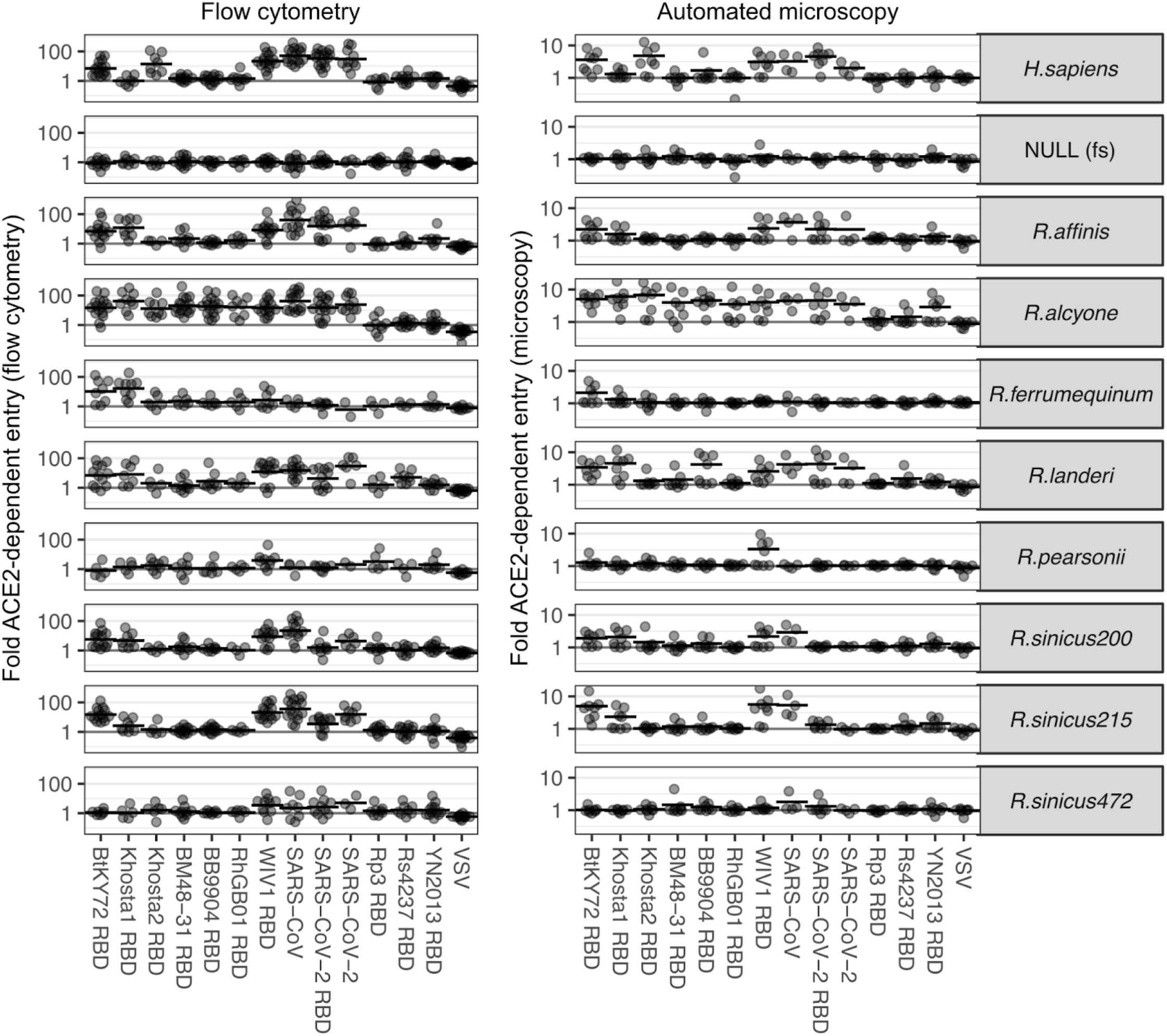
ACE2-dependent infection of various chimeric RBD spike pseudoviruses and cells overexpressing various ACE2 orthologs. ACE2-dependence values calculated following flow cytometry values are shown on the left, while values calculated through microscopy are shown on the right. All experiments were performed with HEK 293T cells overexpressing the indicated ACE2 sequence and human TMPRSS2 co-translationally linked together with a 2A translational stop-start sequence.

**SFig 6.**
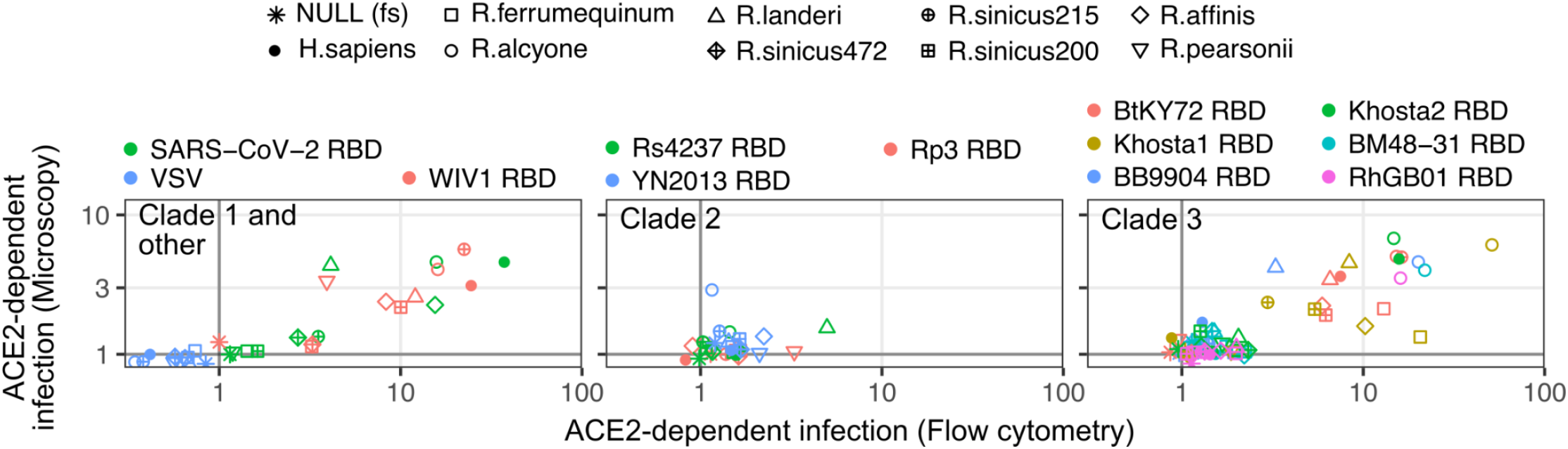
Correlations between flow cytometry and microscopy infection readouts. Scatter plots showing the correlation of ACE2-dependent infection quantitated with flow cytometry (x axis) and microscopy (y axis), separated by sarbecovirus clades. All experiments were performed with HEK 293T cells overexpressing both ACE2 and human TMPRSS2, linked together with a 2A translational stop-start element.

**SFig 7.**
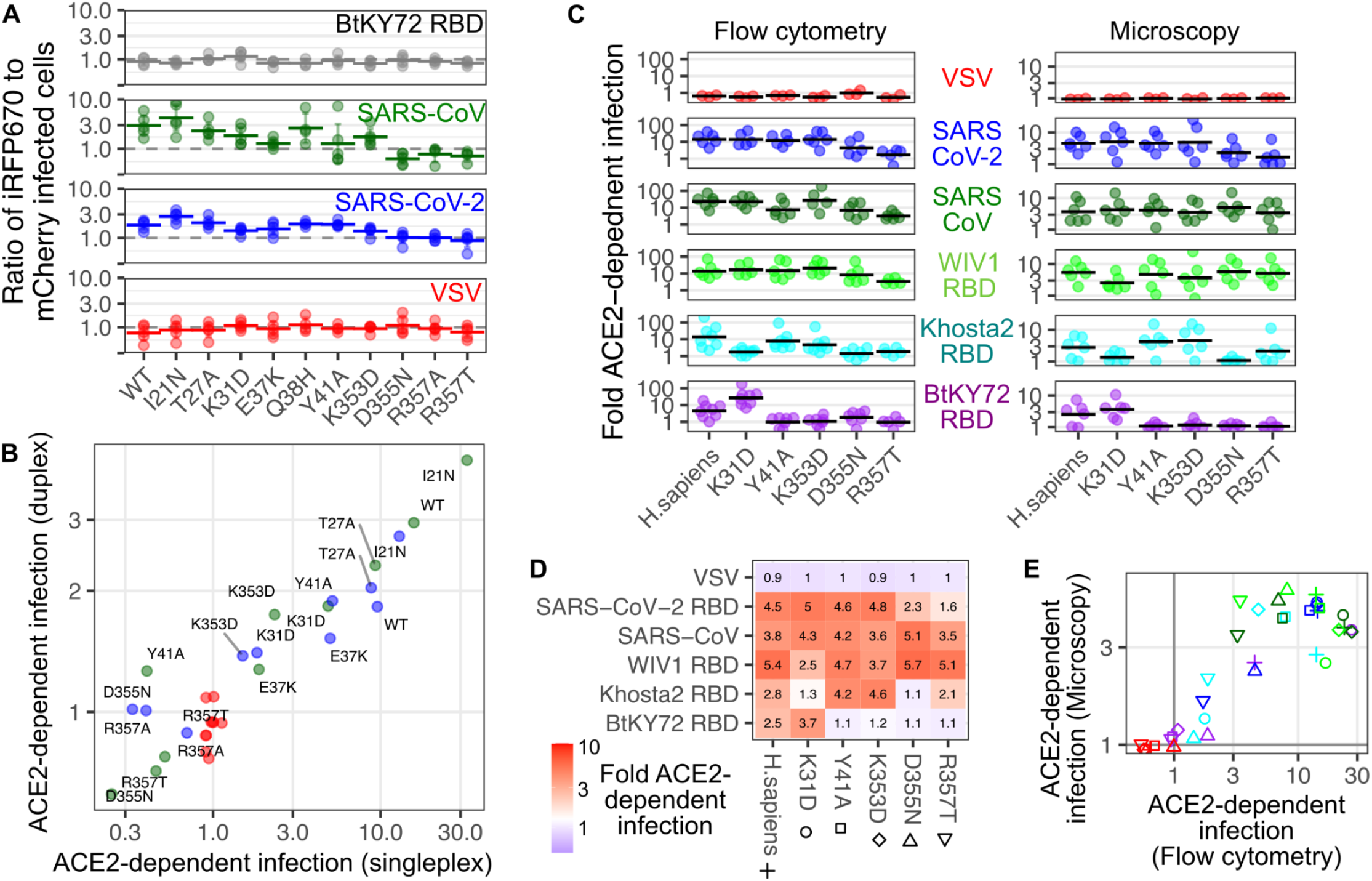
ACE2 amino acid requirements observed with clade 3 sarbecovirus RBDs. A) ACE2-dependent infection of cells expressing WT or variant human ACE2 with chimeric RBD pseudoviruses. These cells expressed relatively low cell-surface ACE2 due to translation of the transgenic mRNA stimulated by a suboptimal Kozak sequence. B) Correlation in ACE2-dependent infectivities as captured by the duplex assay shown in panel A compared with the results we previously obtained using the singleplex assay in a prior publication. C) Flow cytometry and microscopy -based ACE2-dependent values observed with the WT or variant ACE2 proteins when encoded behind a consensus Kozak sequence and coexpressed with TMPRSS2. D) Heatmap showing the ACE2-dependent values obtained through microscopy. E) Correlation of geometric mean ACE2-dependent infectivities observed with the flow cytometry and microscopy readouts of the infection assay. Colors are as shown in panel C. Symbols are as labeled in panel D.

**SFig 8.**
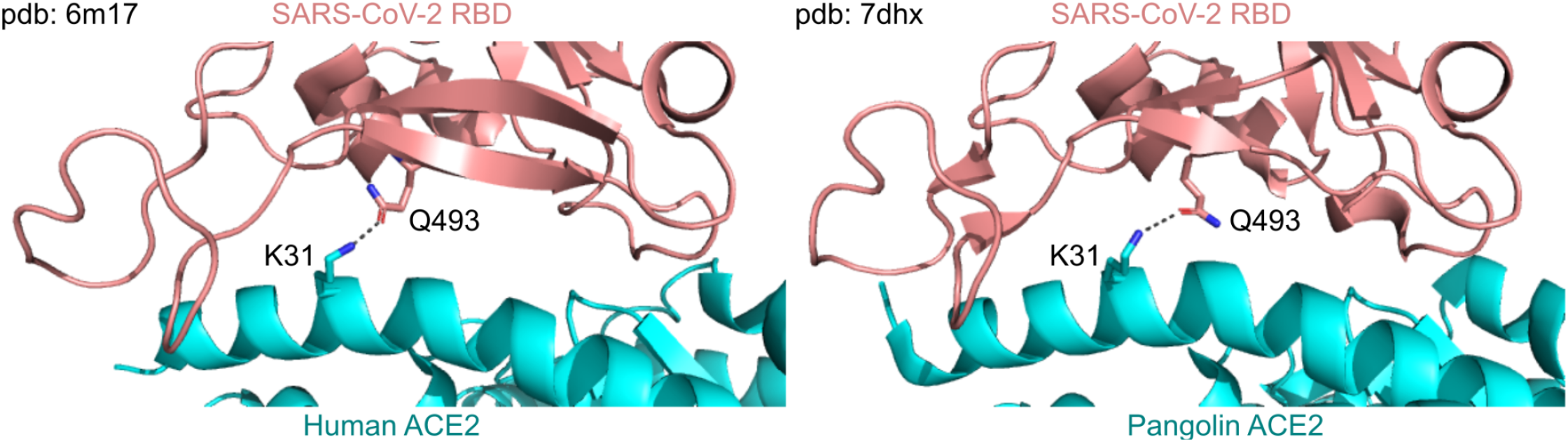
Protein structures with hydrogen bonds between ACE2 K31 and SARS-CoV-2 RBD Q493. A) A cryo-electron microscopy structure (PDB: 6m17, left) and an X-ray diffraction structure (PDB: 7dhx, right) between SARS-CoV-2 RBD and either human (left) or pangolin (right) ACE2. The side chain residues for ACE2 Lys31 and SARS-CoV-2 RBD Gln493 are shown as stick representations, with nitrogen atoms colored blue and oxygen atoms colored red. ACE2 is colored cyan, and SARS-CoV-2 RBD is colored salmon. The predicted hydrogen bond is shown as black dashes.

## Notes

**Funding**: This research was supported by a National Institutes of Health (NIH) grant AI141620 (KAM), AI156907 (KAM), GM142886 (KAM), and AI161275 (AMB). The Cytometry & Imaging Microscopy Shared Resource of the Case Comprehensive Cancer Center was supported by NIH grants P30CA043703 and S10OD021559.

**Competing interests**: None

### Competing Interest Statement

The authors have declared no competing interest.

